# Integrating affinity chromatography in the platform process for Adenovirus purification

**DOI:** 10.1101/2025.03.31.646491

**Authors:** Yuxuan Wu, Eduardo Barbieri, William K. Smith, Arianna Minzoni, Ryan E. Kilgore, Wenning Chu, Michael A. Daniele, Stefano Menegatti

## Abstract

Adenoviral vectors (AdVs) are gaining prominence in cancer therapy and vaccine development, posing the need for a modern AdV manufacturing platform. Current AdV purification by ion-exchange chromatography indeed struggles to achieve the product’s yield and purity of processes that employ affinity technologies. Addressing these challenges, this study presents the first affinity-based process that delivers high product yield and clearance of host cell proteins and DNA (HCPs and hcDNA) in two chromatography steps. The affinity capture utilizes resins functionalized with peptide ligands that target AdV hexon proteins (AEFFIWNA and TNDGPDYSSPLTGSG), and provide high capacity (>5·10^10^ vp per mL of resin) and yield under mild elution conditions (∼50% at pH 8.0). Peptide-functionalized adsorbents prepared using different matrices (polymethylmethacrylate *vs*. agarose) were initially tested to compare the purification performance. AEFFIWNA-SulfoLink™ resin was selected for its yield of cell- transducing AdVs (∼50%) and removal of HCPs and hcDNA (144-fold and 56-fold). Similarly, TNDGPDYSSPLTGSG-Toyopearl^®^ resin afforded ∼50% yield and >50-fold reduction of contaminants. Additional gains in product purity were achieved by optimizing the washing step, which removed free hexon proteins and additional HCPs. All peptide-functionalized resins maintained their purification performance for ten cycles upon regeneration at pH ∼2.0. The purification process was assembled to include clarification, affinity capture in bind-and-elute mode using AEFFIWNA-SulfoLink™ resin, and polishing in flow-through mode using mixed-mode resins. The optimized process provided a yield ∼50% of cell-infecting units (IFU) and a product titer ∼10^7^ IFU/mL, along with residual HCP and hcDNA levels (∼10 ng/mL and 44 ng per dose, respectively) that meet clinical requirements.

## 1. Introduction

The adoption of Adenoviral vectors (AdVs) for delivering mRNA vaccines during the Covid19 pandemic has brought AdV back in the spotlight of the biomanufacturing community. Since 2020, the clinical trials employing AdVs for cancer therapy (tumor suppressor and suicide genes, and immunostimulatory genes) (Barton et al., 2006; Gao et al., 2020; Sorensen et al., 2009; Wang et al., 2009), vaccine development (Afkhami et al., 2016; Sakurai et al., 2022; Thacker et al., 2009), and regenerative therapies and stem cell engineering (Barkats et al., 1998; Tasca et al., 2020) have grown steadily reaching over ninety registered entries in 2024, a level second only to the gene therapy trials of adeno-associated viruses (AAVs) (Lee et al., 2017). Unlike AAVs, AdVs can deliver genes of significantly larger size (up to 36 kbases) and their broad range of serotypes (more than 60 human AdVs have been identified to date) enables the selective infection of various human cells with high transfection efficiency (Nadeau and Kamen, 2003; Ricobaraza et al., 2020; Syyam et al., 2022; Watanabe et al., 2021). The rising number of pre-clinical and clinical studies, coupled with the anticipated regulatory approvals of AdV-based therapies (the first oncolytic AdV, Adstiladrin, was approved in 2022 for the treating invasive bladder cancer (Anon, 2022; Colbert et al., 2025)) underscores the need for robust and affordable expression and purification technologies.

A notable difference between the processes for the purification of AAVs and AdVs for *in vivo* gene therapies is the presence of an affinity-based capture step. Current AdV purification platform processes, in fact, rely on conventional anion-exchange and hydrophobic interaction chromatography (Brument et al., 2002; Burova and Ioffe, 2005; Ehrke-Schulz et al., 2016; Kaludov et al., 2002; Shabram et al., 1997; Ugai et al., 2005). While capable of achieving the product titer and quality needed for vaccine applications (10^11-12^ vg and < 10 ng of hcDNA per dose), these processes struggle to meet the standards required by oncolytic applications (up to 10^13^ vg, < 100 ng of HCPs, and < 10 ng of hcDNA per dose) (Nokisalmi et al., 2010; Salauddin et al., 2024; Turnbull,; Wold and Toth, 2014; Yang, 2013). An affinity resin for AdV purification, Poros™ CaptureSelect™ AdV5 affinity matrix (Dietl et al., 2023; Gast et al., 2019; Gast et al., 2020), is available but it has not yet been integrated in a full purification process reported in either peer-reviewed or patent literature.

In prior work, our team introduced an ensemble of peptide ligands that selectively target the hexon protein, the most abundant component of the adenoviral capsid (Wu et al., 2024). Specifically, ligand AEFFIWNA was identified by combinatorial screening while ligand TNDGPDYSSPLTGSG was designed *in silico* as a mimetic of anti-hexon antibodies. The hexon targeting selectivity of both ligands was demonstrated by surface plasmon resonance and molecular dynamics simulations. Furthermore, the initial characterization of peptide-functionalized resins demonstrated the ability of these ligands to purify Adenovirus serotype 5 (AdV5) from a HEK 293 and Vero cell lysates achieving ∼50% yield and ∼100-fold reduction of HCPs.

Building on this foundation, we resolved to utilize peptide-functionalized adsorbents to build an affinity-based platform process of AdV purification that achieves superior product yield and purity compared to current processes. To this end, we initially conjugated the peptide ligands on polymethylmethacrylate and crosslinked agarose beads, and selected AEFFIWNA- Sulfolink™ and TNDGPDYSSPLTGSG-Toyopearl^®^ resins for their high binding capacity, product yield and purity obtained upon mild elution conditions, and lifetime upon harsh regeneration conditions. We then integrated AEFFIWNA-Sulfolink™ in a 3-step purification process including clarification, affinity-based capture in bind-and-elute mode, and mixed-mode polishing in flow-through mode. The optimized process featured global yield ∼50% of HEK293 infection dose units (IFUs) and a 500-fold and 10^4^-fold reduction of hcDNA and HCPs, delivering a product titer of ∼10^7^ IFU/mL and residual hcDNA of 44 ng per dose.

## 2. Materials and Methods

### 2.1. Materials

The Toyopearl^®^ AF-Amino-650M resin (pore size: 100 nm; particle size: 65 μm; ligand density: 100 μmol per mL resin) was obtained from Tosoh Corporation (Tokyo, Japan), while SulfoLink™ coupling resin and POROS™ CaptureSelect™ AdV5 affinity matrix were obtained from ThermoFisher Scientific (Waltham, MA, USA). All the Fmoc/tBu- protected amino acids, the hexafluorophosphate azabenzotriazole tetramethyl uronium (HATU) coupling reagent, piperidine, diisopropylethylamine (DIPEA), and trifluoroacetic acid (TFA) were obtained from ChemImpex International (Wood Dale, IL, USA). Peptides AEFFIWNAC and TNDGPDYSSPLTGSGC were purchased from GenScript (Piscataway, NJ, USA). Triisopropylsilane, 1,2-ethanedithiol (EDT), polybrene, phosphate buffered saline at pH 7.4 (PBS), hydrochloric acid (HCl), sodium hydroxide (NaOH), sodium chloride (NaCl), and sodium citrate monohydrate were obtained from MilliporeSigma (St. Louis, MA, USA). N-methyl-2-pyrrolidone (NMP), N,N’-dimethylformamide (DMF), dichloromethane (DCM), methanol, and Bis-Tris HCl were obtained from Fisher Chemical (Hampton, NH, USA). Dulbecco’s Modified Eagle Medium (DMEM), fetal bovine serum (FBS), TaqMan™ Fast Advanced Master Mix, Custom Taqman™ assay GFP probe, Proteinase K, Turbo DNase, DNase buffer, Gibco Viral Production cells HEK293F, Picogreen DNA testing kit were sourced from ThermoFisher Scientific (Waltham, MA, USA). Standard Adenovirus serotype 5 (AdV5) loaded with a transgene encoding for green fluorescence protein was purchased from Vector Biolabs (Malvern, PA, USA). BalanCD HEK293 cell culture medium was obtained from Irvine Scientific (Santa Ana, CA, USA). African Green Monkey Kidney (Vero) cells were obtained from ATCC (Manassas, VA, USA). All chromatographic experiments were performed using an ÄKTA Avant system (Cytiva, Marlborough, MA, USA). Alltech chromatography columns (L/ID: 50/3.6 mm; volume: 0.5 mL), and 10 μm polyethylene frits were obtained from VWR International (Radnor, PA, USA). The Tricorn 5/50 empty chromatography column was obtained from Cytiva (Marlborough, MA, USA). The HEK293 HCP and Vero HCP ELISA kits were purchased from Cygnus (Southport, NC, USA). The SurePAGE™ Bis-Tris Gel 4-12% was bought from GenScript (Piscataway, NJ, USA). The PageRuler™ Plus Prestained Protein Ladder, 10 to 250 kDa was purchased from ThermoFisher Scientific (Waltham, MA, USA). The 4x Laemmli Sample Buffer was purchased from Bio-Rad laboratories (Hercules, CA, USA).

### 2.2. AdV5 culturing and clarification

HEK293F suspension cells were cultured in BalanCD HEK293 medium to a density of 6·10^6^ cells/mL, as measured using an Invitrogen Countess 3 Automated Cell Counter (ThermoFisher Scientific, Waltham, MA). The cells were diluted in BalanCD HEK293 medium to a concentration of 10^6^ cells/mL, incubated with AdV5 at a multiplicity of infection (MOI) of 6.7, and cultured in a thermostatted shaking incubator at 37°C and 8% CO_2_ for 40 hrs. Infected cells were centrifuged 15 mins at 1940 rcf and the supernatant was removed. The pelleted cells were resuspended in 2 mL of 10 mM Tris-HCl at pH 8.0 and lysed via three cycles of freezing at -80°C and thawing at 37°C. The cell debris were removed either by centrifugation at 1940 rcf and 4°C for 15 mins, the crude product was filtered by 0.2μm Fisherbrand™ Disposable PES Filter Units.

### 2.3. Peptide conjugation on different matrix

The hydroxyl groups on Poros™ 50 OH resin were initially converted to primary amines as follows: a volume of 10 mL of resin was initially dried using a stream of nitrogen, washed in DMF, and resuspended in 50 mL of a solution of CDI at 100 mg/mL in DMF; after 5 hours, the resin was copiously washed with DMF and dried with a stream of nitrogen; the imidazole ester-activated resin was then mixed with 100 mL of 5% v/v ethylenediamine in DMF and incubated at 45ºC under shaking at 100 rpm; after 12 hours, the resin was washed with DMF and DCM, dried with nitrogen, and stored at 4ºC. The peptide sequences AEFFIWNA and TNDGPDYSSPLTGSG were then conjugated on Toyopearl^®^ AF-Amino-650M resin and aminated Poros™ 50 via Fmoc/tBu chemistry using an Initiator+ Alstra (Biotage, Uppsala, Sweden). Peptide ligands AEFFIWNAC and TNDGPDYSSPLT-GSGC were conjugated on SulfoLink™ coupling resin following the manufacturer’s instructions. The resins were individually packed in Alltech chromatography columns (L/ID: 50/3.6 mm; volume: 0.5 mL), washed with 20% v/v ethanol, and equilibrated with PBS at pH 7.4.

### 2.4. Purification of AdV5 from HEK293 and Vero cell lysates using peptide-functionalized resins

A volume of 5 mL (10 column volumes, CVs) of cell lysate (AdV titer ∼10^9^ vp/mL; HEK293 or Vero HCP titer ∼ 0.1 mg/mL) was loaded on the column in down-flow at the flow rate of 0.143 mL/min (residence time (RT) of 3.5 min). Following a washing step with 10 CVs of equilibration buffer at the RT of 3.5 min, the bound AdVs were eluted in up-flow using 20 CVs of 1 M NaCl in 20 mM Tris HCl buffer at pH 8.0 at the RT of 3.5 min. Finally, the column was regenerated with 10 CVs of either 10% v/v phosphoric acid at the RT of 3.5 min or 0.1 M glycine buffer at pH 2.0 at the RT of 2 min.

### 2.5. Dynamic AdV5 binding capacity of peptide-functionalized resins

AEFFIWNA- Toyopearl, AEFFIWNAC-Sulfolink, and TNDGPDYSSPLT-GSG-Toyopearl^®^ resins were individually packed into 0.5 mL Alltech columns, washed with 20% v/v ethanol, and equilibrated with PBS at pH 7.4. A volume of 80 CVs of HEK293 cell lysate (AdV titer ∼10^8^ or 10^9^ vp/mL; HCP titer ∼ 0.15 mg/mL) was loaded at the RT of either 2 min or 3.5 min. The effluents were continuously monitored via UV spectroscopy at 260 and 280 nm, and apportioned in fractions of 2 CVs. The adenoviral genome in the collected fractions was measured via qPCR as described in *Section 2.7* and utilized to calculate the dynamic transgene-loaded AdV5 binding capacity at 10% breakthrough (DBC_10%_).

### 2.6. Quantification of HEK293 and Vero HCPs and hcDNA

The titer of HCPs in the feedstocks and chromatographic fractions was measured using HEK293 and Vero HCP ELISA kits (Cygnus Technologies, Southport, NC) following the manufacturer’s protocols. The quantification of host cell DNA (hcDNA) was performed using Quant-iT™ PicoGreen™ dsDNA Assay Kits (ThermoFisher Scientific, Waltham, MA) following the manufacture’s protocol. The values of HCP and hcDNA logarithmic reduction values (LRV) were calculated by mass balance.

### 2.7. Quantification of adenoviral genome

The titer of AdV-encapsidated transgenes in the feedstocks and chromatographic fractions was measured via qPCR (*note:* ELISA kits for the quantification of AdV particles are not commercially available). The samples were initially treated with DNase to remove free viral and host cell DNA: briefly, 20 μL of sample were mixed with 2 μL Turbo DNase buffer and 1 μL DNase, and incubated at 60°C for 60 mins and then 95°C for 20 mins. Thereafter, 1 μL of Proteinase K was added to each sample, followed by incubation at 37°C for 60 mins and then at 95°C for 10 mins. qPCR was then conducted in a CFX Duet real time PCR system (Biorad Laboratories, Hercules, CA) using TaqMan probes (forward primer sequence: AGCAAAGACCCCAACGAGAA; reverse primer sequence: GGCGGCGGTCACGAA; GFP probe sequence: CGCGATCACATGGTCCTGCTGG) to quantify the GFP gene.

### 2.8. Quantification of adenovirus recovery by median tissue culture infectious dose (TCID50)

Aliquots of 100 μL of a suspension of HEK293 at the cell density of 10^5^ cells/mL in DMEM-2 were dispensed in 96-well plates. A 12-fold serial dilution of an AdV5 feedstock was prepared encompassing the titer range of 10^9^ to 10^10^ vg/mL. Aliquots of 100 μL of each dilution were added to the 96 well plate, with 9 replicates per titer, while 100 μL of DMEM-2 medium were used as controls. The 96 well plates were incubated at 37°C for 10 days and then analyzed to detect cytopathic effects (CPE, any morphological change detected in a well was logged as positive). The CPE ratio was calculated for every one of the 10 values of dilution (10^−3^, 10^−4^, …, 10^−11^) as the number of cells in which CPE was detected divided by the total number of cells (herein, 9). The resulting 10 values of CPE ratios were added to obtain the score “S”. The final TCID50 value was calculated according to the formula T = 10 (3+S) (Dietl et al., 2023; Iyer et al., 1999; Wu et al., 2002).

### 2.9. Electrophoretic analysis of the chromatographic fractions

SDS-PAGE analysis of the feedstocks and the fractions obtained by purifying AdV5 using AEFFIWNA-functionalized resins was conducted using 4-12% SurePage™ gels (Genscript, NJ) and 1X Tris-MOPS-SDS Running Buffer (Genscript, NJ) as running buffer. The gel was run on a Mini-PROTEIN Tetra cell system (Bio Rad, CA). A volume of 30 μL of sample was combined with 10 μL of 4x Laemmli Sample Buffer and loaded to the wells of SDS-PAGE gels. The gel was run at 100 V for about 10 min and 200 V for about 30 minutes. The gels were then stained using a SilverQuest™ Silver Staining Kit (ThermoFisher, Waltham, MA) and finally imagined under FBWLT-1417 White Light Transluminator (FisherBiotech) and extracted with ImageJ v1.54 (Chu et al., 2022; Chu et al., 2023).

### 2.10. Analysis of the chromatographic fractions size-exclusion chromatography (SEC)

The feedstock and chromatographic fractions collected as described in *Section 2.4* were analyzed by SEC-HPLC using a BioResolve SEC mAb Column (200Å, 2.5 μm, 4.6 × 300 mm; Waters, Milford, MA) operated in isocratic mode using 200 mM KCl in 50 mM sodium phosphate at pH 7.0 (0.05% v/v sodium azide) as mobile phase, and at the flow rate of 0.5 mL/min for 40 min or a Bio SEC-5 HPLC column (2000 Å, 5 μm, 4.6 × 300 mm; Agilent Santa Clara, CA) operated in isocratic mode using the same mobile phase at 0.4 mL/min for 30 min. A volume of 20 μL of sample was injected and the effluent was continuously monitored via UV (260 and 280 nm) fluorescence spectroscopy (ex/em: 280/350 nm) (Chu et al., 2023).

## 3. Results

### 3.1. Purification of AdV5 from HEK293 and Vero cell lysates using peptide-functionalized resins under optimal elution conditions

Our initial study introducing peptide ligands AEFFIWNA and TNDGPDYSSPLTGSG presented the purification of Adenovirus serotype 5 (AdV5) from HEK293 cell lysates as a demonstrative case study(Wu et al., 2024). Our initial evaluation focused on quantifying the yield of encapsulated transgenes (*i.e*., reporter gene encoding green fluorescent protein, GFP) and the removal of host cell proteins (HCPs). In this investigation, we seek to broaden the characterization of these ligands by adding the measurement of yield of cell-transducing AdV5 particles and the clearance of host cell DNA (hcDNA) under optimized elution conditions. To this end, we employed clarified cell lysates featuring an AdV5 titer ∼10^9^ vg/mL and HCP titers ranging from ∼0.08-0.35 (HEK293F cells) to ∼0.1 mg/mL (Vero cells) as shown in **Table S1**. The feedstocks were loaded on columns packed with 0.5 mL of AEFFIWNA-Toyopearl^®^ and TNDGPDYSSPLTGSG-Toyopearl^®^ resins at ratios of 10^9^∼10^10^ vg per mL of resin and a residence time (RT) of 3.5 min. The resulting values of yield of encapsidated transgenes and infectious units (IFU), and the corresponding logarithmic removal values of host cell contaminants (HCPs and hcDNA LRVs) are presented in **Figure 1** and **Table S1**.

**Figure 1.**
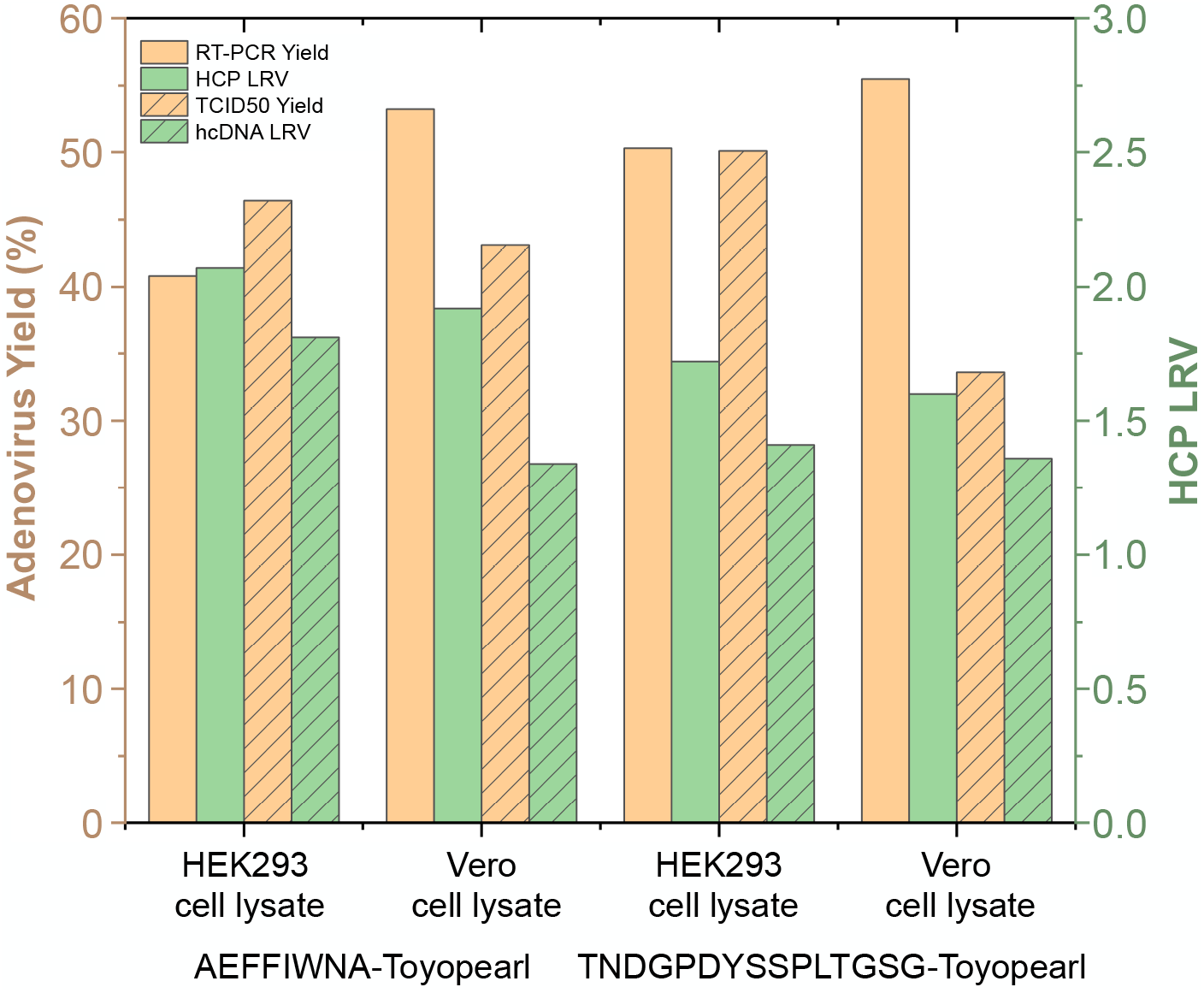
Performance of AdV5 purification from clarified HEK293 and Vero cell lysates (AdV5 titer ∼10^9^ vg/mL; HCP titer ∼0.08 – 0.35 mg/mL) using AEFFIWNA-Toyopearl® and TNDGPDYSSPLTGSG-Toyopearl® resins. The values of AdV5 yield were measured via RT-qPCR (encapsidated transgenes) and TCID50 assays (cell-transducing units); the reduction of HCPs and hcDNA were measured by analyzing the eluates and corresponding feedstocks using ELISA and PicoGreen™ dsDNA assay kits.

AEFFIWNA-Toyopearl^®^ resin achieved AdV5 transgene yields of 40.5% and 53.2% from the HEK293 and the Vero cell lysates, respectively, along with average reduction values of ∼100-fold for HCPs and 60-fold for hcDNA; TNDGPDYSSPLTGSG-Toyopearl^®^ resin afforded slightly higher yields, respectively 50.3% and 55.5%, but lower clearance of HCPs (50- fold) and hcDNA (30-fold). It is noted, however, that the different reduction values stem from the differences in the titers of contaminants in the HEK293 *vs*. Vero cell lysates, whereas the residual titers of HCPs and hcDNA in the eluates are similar, namely 0.9 – 1.5 μg/mL and 120 – 150 ng/mL (*note:* the removal of hcDNA could be further improved by a DNase treatment after cell lysis). For reference, AdV purification by anion exchange offers higher recoveries (up to 80%) but significantly lower removal of host cell contaminants (Ehrke-Schulz et al., 2016). Furthermore, the use of anion exchange requires conducting a UF/DF step prior to product capture to increase the AdV titer and reduce the conductivity and level of contaminants in the clarified cell lysate.

When operated with the elution conditions identified in prior work (1 M NaCl in 20 mM Tris buffer at pH 8.0) (Wu et al., 2024), the peptide ligands provided high yields of encapsidated transgenes and cell-transducing virus particles from both cell lysates, showing negligible product loss despite the high loading (<1% of loaded AdV5 was lost during the loading and washing steps), and efficient removal of host cell contaminants. However, since the individual roles of conductivity and pH on the AdV5:peptide interaction were unknown, we resolved to investigate the dependence of product yield on the formulation of the elution buffer. To this end, we implemented a step-wise increase in NaCl concentration from 0.2 M to 1.2 M (*i.e*., four 0.2 M NaCl increments of 5 CVs each), while maintaining a constant pH 8.0. The results in **Figure 2A** show that ligand AEFFIWNA released small amounts (< 3%) of AdV5 at low conductivity, whereas satisfactory product recovery could be achieved only at high conductivity (step yields of ∼10% and ∼30% were recorded at 0.8 M and 1 M NaCl, respectively). Conversely, the release of AdV5 from ligand TNDGPDYSSPLTGSG increased linearly with conductivity, starting from a step yield of 4% at 0.2 M NaCl and reaching 14% at 1 M NaCl. The sum of the step yields reported in **Figure 2A** match the values (40 - 55%) presented in **Figure 1**, suggesting the need to identify orthogonal triggers that dissociate the AdV5:peptide complex and increase product yield.

**Figure 2.**
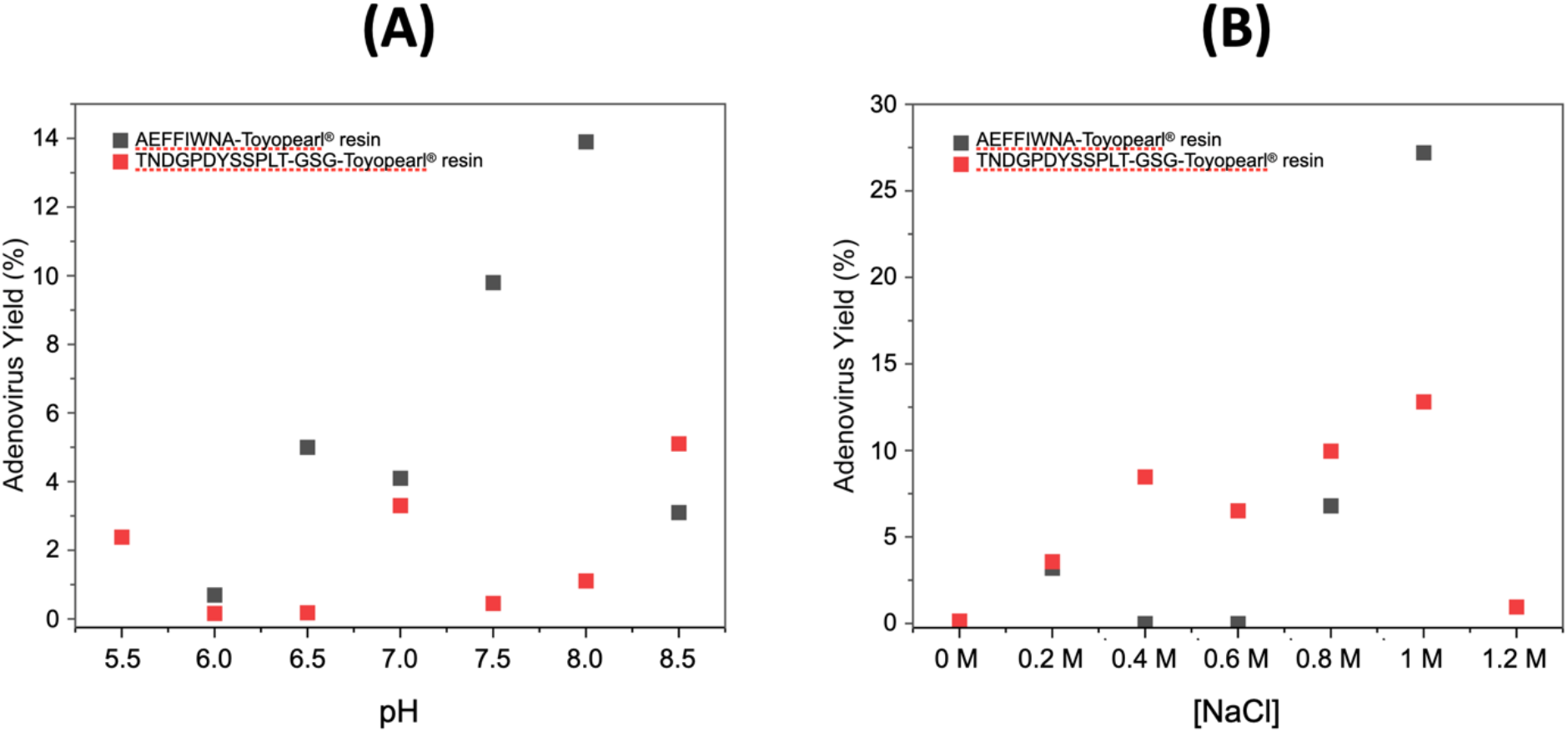
Dependence of yield of AdV5 purified from a clarified HEK293 cell lysate (AdV5 titer ∼10^9^ vg/mL; HCP titer ∼0.15 mg/mL) using AEFFIWNA-Toyopearl® and TNDGPDYSSPLTGSG-Toyopearl® resins upon **(A)** NaCl concentration and **(B)** pH of the elution buffer. The values of AdV5 yield were measured via RT-qPCR (encapsidated transgenes) analysis of the eluates and corresponding feedstocks.

The optimization of the elution pH was conducted within the range 6.0 – 8.5, while fixing the NaCl concentration at 1 M. At pH above 8.5, AdV capsids undergo structural changes that lead to a loss of viral infectivity. Similarly, lowering the pH to below 6 triggers a premature disassembly of the capsid, specifically release of vertex proteins (Wiethoff et al., 2005). Although it has been reported that activity losses can be partially reverted when the lower pH is returned to physiological values (Wold and Toth, 2014), we set the lower and upper limits of the elution pH gradient to 6.0 and 8.5. As shown in **Figure 2B**, the pH dependency of AdV5 release from AEFFIWNA followed the same profile observed with conductivity, where only a small amount (2 - 5%) of virus was released between pH 6.0 and 7.0, followed by an uptick to ∼10 - 15% at pH 7.5 and 8.0. Conversely, the step yield of AdV5 from TNDGPDYSSPLTGSG featured a bi-modal profile, with peaks of 3% and 5% at pH 7.0 and 8.5, while being near null at all other values.

These results match the outcomes of the molecular docking and dynamics simulations conducted in prior work, showing that the AdV5:AEFFIWNA interaction at pH 7.4 is dominated by hydrogen bonds, while hydrophobic and coulombic interactions only play an ancillary role. Accordingly, the addition of NaCl, a mild kosmotrope, is expected to disrupt the hydrogen bonding network and promote the formation of a water shell around the capsid, thereby inducing the dissociation of the AdV:peptide complexes. Additionally, as the pH increases to 8.0 – 8.5, the repulsive effect of the ligands’ anionic residues Asp (D) and Glu (E) to their counterparts on the capsid furthers the release of AdV5. Accordingly, PBS at pH 7.4 and 1 M NaCl in 20 mM Tris buffer at pH 8.0 were maintained respectively as binding and elution buffers. This choice is advantageous since these formulations are already utilized for adenovirus purification via anion exchange chromatography, and enables as seamless integration of the peptide-functionalized adsorbents in current bioprocesses.

### 3.2. Matrix selection of target sequence

The design of chromatographic adsorbents – chiefly, the composition of the beads, their particle and pore diameters, and the ligand density – is key to optimal binding capacity, selectivity, and product yield. While extensive research exists on the composition and morphology of affinity and ion-exchange resins for the purification of proteins (≤ 10 nm), much less is known on their role in purifying complex biologics. Viral vectors in particular are challenging due to their larger size (20 - 100 nm), biomolecular complexity of their surface, and diverse product-related impurities (*e.g*., capsid proteins and capsomers, and full *vs*. empty capsids). Driven by the growing importance of gene therapies, bioseparation researchers have begun to prioritize the development of chromatographic substrates tailored to gene therapies, focusing primarily on adeno-associated viral vectors (AAVs) and plasmids/mRNA. However, similar efforts on lentiviral and adenoviral vectors (LVVs and AdVs ∼80-100 nm) are lagging. To address this gap, this study investigates the roles of chromatographic substrate composition and morphology on AdV affinity purification by comparing *(i)* polystyrene-based beads with large pores (Poros™, particle size: 50 μm, pore size: 100-1000 nm), the same matrix utilized in the commercial CaptureSelect™ AdV5 affinity resin (Gast et al., 2019; Gast et al., 2020); *(ii)* poly(methyl methacrylate)-based beads with medium pores (ToyoPearl^®^, 65 μm, 100 nm), selected for its high rigid pore structure and ability to operate at high flow rates; and *(iii)* crosslinked agarose resins, selected for its high hydrophilicity that grants minimal non-selective adsorption (Chu et al., 2022; Chu et al., 2023). The purification performance of the resulting affinity adsorbents functionalized with peptides AEFFIWNA and TNDGPDYSSPLTGSG is presented in **Table S2** and summarized in **Figure 3** (the corresponding chromatograms are collated in **Figure S3**).

**Figure 3.**
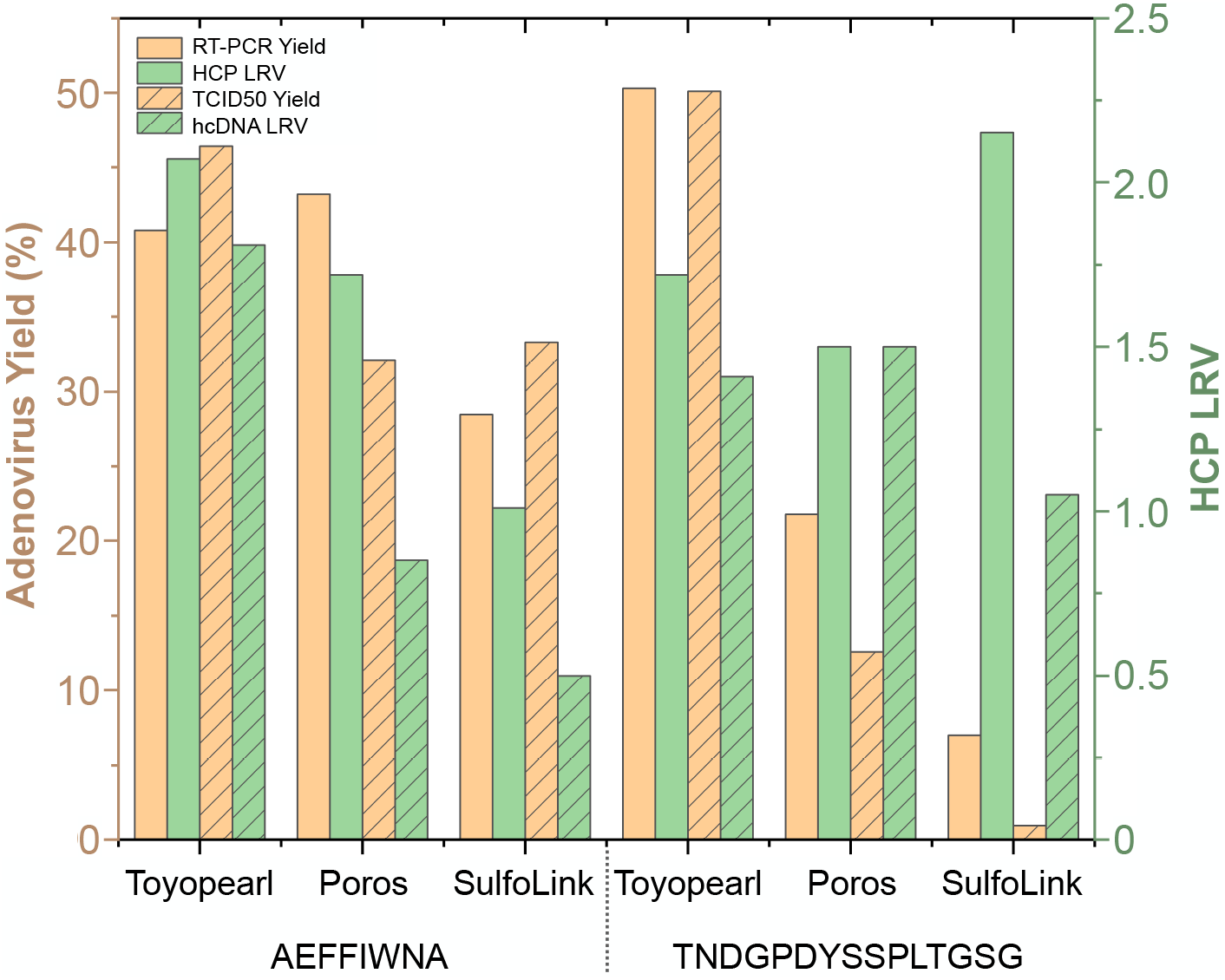
Performance of AdV5 purification from a clarified HEK293 cell lysate (AdV5 titer ∼10^9^ vg/mL; HCP titer ∼0.15 mg/mL) using peptide ligands AEFFIWNA and TNDGPDYSSPLTGSG conjugated to Poros™, Toyopearl®, or SulfoLink™ resins. The values of AdV5 yield were measured via RT-qPCR (encapsidated transgenes) and TCID50 assays (cell-transducing units); the reduction of HCPs and hcDNA were measured by analyzing the eluates and corresponding feedstocks using ELISA and PicoGreen™ dsDNA assay kits.

The AdV5 capture and release recorded on AEFFIWNA-functionalized resins are comparable for Toyopearl^®^ and Poros™ beads, which eluted 6.2·10^9^ and 4.0·10^9^ vg per mL of resin (corresponding to yields of 40.8% and 43.2%, respectively). Conversely, the SulfoLink™ resin loaded and released up to 5.6·10^10^ and 8.0·10^9^ vg/mL, corresponding to a remarkable productivity of 8.0·10^10^ vg per mL of resin. In comparison, the productivities of AdV5 purification by CaptureSelect™ AdV5 affinity resin and SepharoseQ anion exchange resin are 2.66·10^10^ and 3.4·10^12^ vg per mL of resin (Inger Salomonsson, 2022; Wu et al., 2024). The larger pore diameter of polymer resin, especially Poros™ beads, translates in a lower specific surface and hence a lower binding capacity. Additionally, larger pores facilitate a deeper penetration of virus particles towards the bead’s core during the loading step (residence time, RT: 3.5 min), which may prove disadvantageous during the elution, as the virions may not have adequate time to diffuse out from the beads. On the front of product purity, AEFFIWNA-Toyopearl^®^ beads afforded the highest reduction values of HCPs (117-fold) and hcDNA (65-fold). Conversely, AEFFIWNA- SulfoLink™ resin afforded an unsatisfactory clearance of host cell contaminants, which countered our expectation of low non-selective adsorption on this hydrophilic, agarose-based matrix. This may be imputed to the SulfoLink™ resin having pores whose size is comparable to the hydrodynamic diameter of the AdV particles (∼100 nm) but being more tortuous than those of Toyopearl^®^ resins.^[16]^

Ligand TNDGPDYSSPLTGSG demonstrated an effective purification performance only on Toyopearl^®^ resin, achieving a yield of 50.3% and a 50-fold removal of host cell contaminants. Conversely, Poros™ and SulfoLink™ resins offered insufficient product yield (21.8% and 7%, respectively) and was therefore abandoned.

### 3.3 Optimizing the AdV5 purification protocol using peptide-functionalized agarose resin

Despite the significant number of active AdV5 particles eluted from AEFFIWNA-SulfoLink™ resin, the value of yield, as the ratio of product recovered relative to the amount loaded, is relatively low (∼27%). Additionally, the poor removal of contaminants is at odds with the inherent hydrophilicity of agarose and the weakly anionic character of its surface. We attributed these issues to the occlusion of resin pores by the AdV5 particles during loading, which results in low yield and the concomitant release of entrapped impurities alongside virions. Despite these challenges, the binding capacity of the adsorbent is remarkable, and the inherent binding selectivity of the ligand can be leveraged to increase product quality. On the other hand, AEFFIWNA-Toyopearl^®^ resin, while providing higher purity, affords a significantly lower binding capacity than anion-exchange resins for AdV purification. We therefore resolved to optimize the design and operation of the AEFFIWNA-functionalized resin to fulfill its potential for the affinity purification of AdV.

To improve the performance of AEFFIWNA-Sulfolink™, we resolved to optimize the chromatographic protocol prior to the elution step by incorporating a secondary wash step and optimizing the residence time during the loading step. After loading the adsorbent with the HEK293F cell lysate (AdV titer of 1.6·10^9^ vg/mL) and conducting an initial wash with PBS at pH 7.4, we applied a step-wise gradient of NaCl from 0.15 to 0.25 M in 50 mM phosphate buffer at pH 7.4, prior to elution with 1 M NaCl in 20 mM Tris buffer at pH 8.0. The analysis of the wash and elution fractions in **Figure 4** indicate that an additional amount of host cell contaminants was removed during the wash step and that monomeric hexon proteins (∼116 kDa) were released as the salt concentration increased up 0.25 M NaCl, yet without suffering huge loss of AdV virions. The analysis of the chromatographic effluents by RT-qPCR (**Figure 4B**) indicated only a small fraction of the encapsidated transgenes loaded on the resin was lost throughout the six wash fractions (5.4% and 3.2% in W1 and W6, respectively). Accordingly, the bands identified in the electrophoretic analysis in **Figure 4A** can be reliably attributed to monomeric capsid proteins or capsomers, rather than full virions. Notably, the analysis of the eluted fraction indicated that the product yield rose to 56.1% and the HCP LRV reached 1.76. Notably, the removal of product-related impurities (*i.e*., empty capsids and capsid proteins) by the additional wash step significantly improves product safety by reducing immunogenic species without affecting the therapeutic efficacy of the dose.

**Figure 4.**
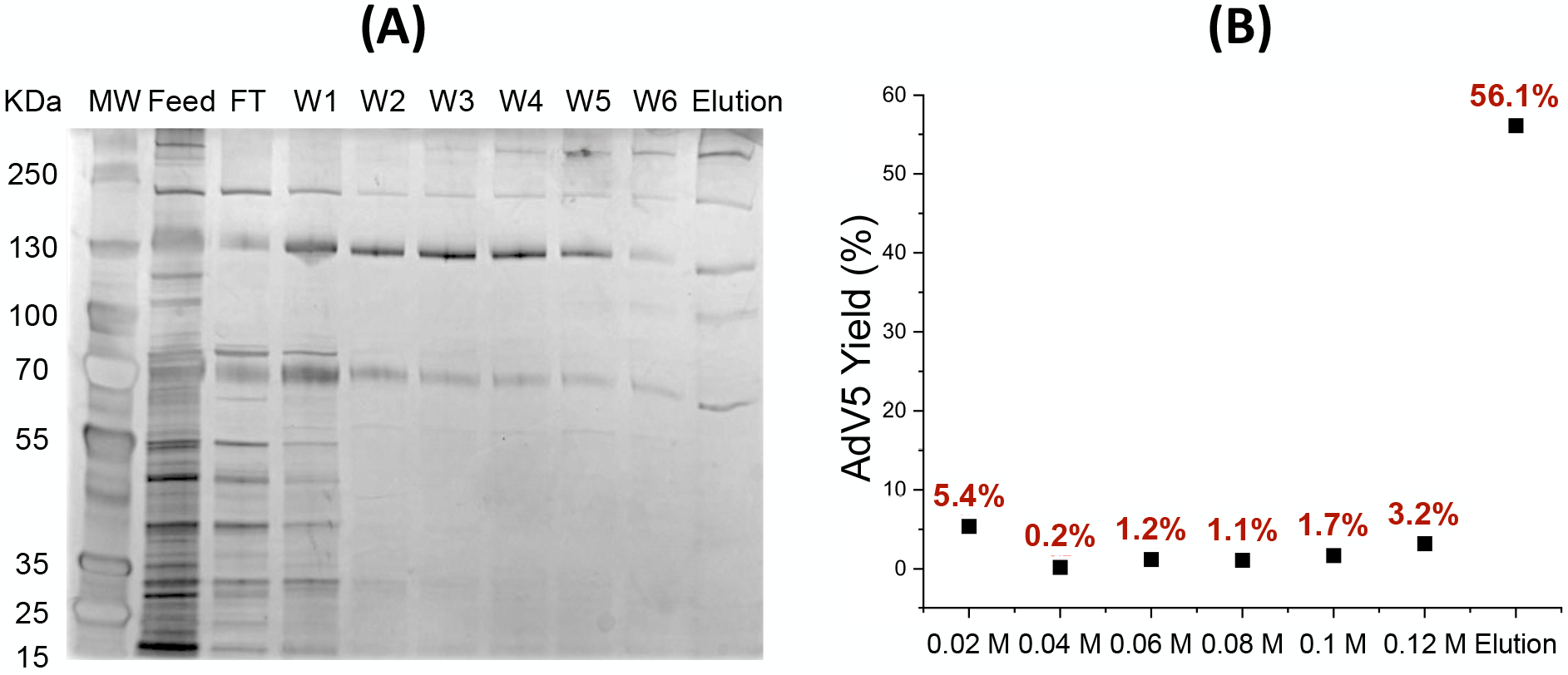
**(A)** SDS-PAGE (non-reducing conditions) analysis of the chromatographic fractions produced by purifying AdV5 from a clarified HEK293 cell lysate (AdV5 titer ∼9*10^9^ vg/mL; HCP titer ∼0.15 mg/mL) using AEFFIWNA- Sulfolink™ resin; lane 1: molecular weight marker; lane 2: feedstock; lane 3: flow-through (5 CVs); lane 4: 1^st^ wash step at 2% elution buffer mixing with loading buffer (5 CVs); lane 5: 2^nd^ wash step at 4% elution buffer mixing with loading buffer (5 CVs); lane 6: 3^rd^ wash step at 6% elution buffer mixing with loading buffer (5 CVs); lane 7: 4^th^ wash step at 8% elution buffer mixing with loading buffer (5 CVs); lane 8: 5^th^ wash step at 10% elution buffer mixing with loading buffer (5 CVs); lane 9: 6^th^ wash step at 12% elution buffer mixing with loading buffer (5 CVs); lane 10: elution (10 CVs); **(B)** values of AdV5 recovery in the wash fractions and in the eluate measured via RT qPCR (encapsidated transgenes).

The results reported above were obtained by loading the feedstock on the columns at the RT of 3.5 min. During the load step, we observed that the AdV5 titer in the effluent (C_FT_) fluctuated significantly (**Figure S2**). We hypothesize that the capture of a large virus using a small linear peptide ligand may lead to a “supersaturated” binding state when the adsorbent is loaded at a small flowrate: during the first phase of loading, the virus saturates the binding sites on the resin; as the load progresses, virus adsorption reaches a supersaturated state; additional virus particles disrupt the supersaturated state, releasing a wave of virus in the effluent (recorded spike in C_FT_) that frees binding sites on the resin’s surface; additional loading allows reaching a second supersaturated state, creating cycles of unstable adsorption and desorption that translate in a fluctuating temporal profile of C_FT_. To test this hypothesis, we increased the loading flow rate, anticipating that a lower contact time – namely, from 3.5 to 2 min – would postpone the formation of a supersaturated state. The breakthrough curves obtained with AEFFIWNA-Toyopearl^®^, and TNDGPDYSSPLTGSG-Toyopearl^®^ resins at the RT of 2 min show conventional profile (**Figure 5**), with C_FT_ rapidly reaching the feedstock titer after saturation. The resultant values of dynamic binding capacities (DBC_10%_), summarized in **Table 1**, consistently indicated that higher binding was achieved at lower residence time, which conducive to higher productivity. Notably, the binding capacity of AEFFIWNA-functionalized resin is higher than that of its commercial counterpart CaptureSelect™ AdV5 affinity resin when operated at higher flow rate, demonstrating the potential of peptide AEFFIWNA as an affinity ligand for the purification of AdV-based therapies.

**Table 1.** Values of dynamic AdV5 binding capacity (DBC_10%_) of AEFFIWNA- and TNDGPDYSSPLTGSG-functionalized resins loaded with a clarified HEK293 cell lysates at different loads and at the RT of either 2 or 3.5 min. The AdV5 titer of 2 min and 3.5 min is the average of two loadings, and there are differences in virus loading concentrations.

| Peptide-functionalized resin | AdV5 titer (vg/mL) | RT | $DBC_{10\%}$ (vg/mL resin) |
| --- | --- | --- | --- |
| AEFFIWNA-Toyopearl® | $\sim 1.7 \cdot 10^9$ | 2 min | $4.6 \cdot 10^{10}$ |
| | | 3.5 min | $2.4 \cdot 10^{10}$ |
| | $3.7 \cdot 10^8$ | 3.5 min | $1.5 \cdot 10^{10}$ |
| AEFFIWNAC-SulfoLink™ | $\sim 1.0 \cdot 10^{10}$ | 2 min | $8.8 \cdot 10^{10}$ |
| | | 3.5 min | $1.3 \cdot 10^{10}$ |
| | $6.3 \cdot 10^8$ | 3.5 min | $9.2 \cdot 10^9$ |
| TNDGPDYSSPLTGSG-Toyopearl® | $\sim 1.7 \cdot 10^9$ | 2 min | $2.3 \cdot 10^{10}$ |
| | | 3.5 min | $1.4 \cdot 10^{10}$ |
| | $2.7 \cdot 10^8$ | 3.5 min | $6.9 \cdot 10^9$ |
| CaptureSelect™ AdV5 | $1.7 \cdot 10^9$ | 3.5 min | $2.8 \cdot 10^{10}$ |

**Figure 5.**
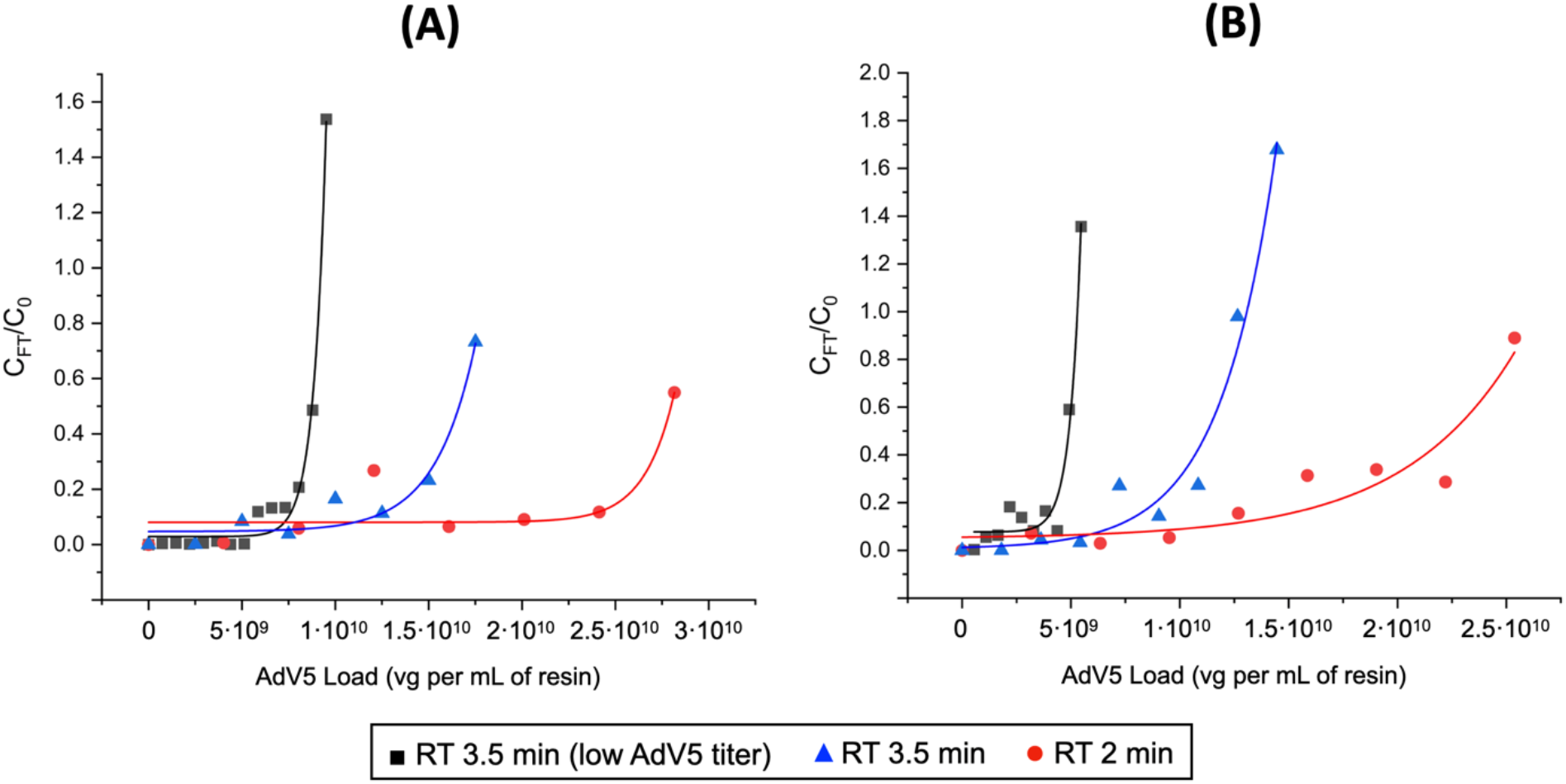
Breakthrough curves of AdV5 encapsidated transgenes obtained by loading clarified HEK293 cell lysates at normal or low AdV5 titers (**Table 1**) on AEFFIWNA- and TNDGPDYSSPLTGSG-Toyopearl® resins loaded with a clarified HEK293 cell lysates at different loads and at the RT of either 2 or 3.5 min.

Guided by the results of DBC_10%_, we proceeded to evaluate the purification performance of AEFFIWNA-Sulfolink™ resin by reducing the loading RT to 2 min and implementing the optimized wash and elution steps (*i.e*., first wash with PBS at pH 7.4; second wash with 0.25 M in 50 mM phosphate buffer at pH 7.4; elution with 1 M NaCl in 20 mM Tris buffer at pH 8.0). The chromatograms are collated in **Figure S4**. The results in **Figure 6A** indicate that product yield grew from 43.2% to 51.2%, in line with the values characteristics of anion exchange chromatography, while the reduction of HCPs and hcDNA rose to 144-fold and 20-fold, respectively. We then proceeded to implement the optimized purification protocol on the AEFFIWNA-SulfoLink™ resin. In preparation to this study, we produced clarified lysates with AdV5 titers of ∼5·10^9^ and ∼2.0·10^10^ vg/mL, which allowed to evaluate the combined effects of loading titer and flow rate on product yield and quality. Notably, the combination of higher feedstock concentration and flow rate shortens the loading phase (*i.e*., the longest step of the chromatographic protocol), thus increasing the productivity of the purification process. Furthermore, loading 10 CVs of feedstock at 5.2·10^9^ vg/mL and 5 CVs of feedstock at ∼2.0·10^10^ vg/mL increased product yield to 51.2% and 61.8%, respectively. In particular, the amount of AdV present in the first 5 CVs of elution increased from 30.7% to 54.8%, indicating that increasing the product titer in the load and reducing the loading time are conducive to increasing both yield and productivity. In sum, the optimal AdV purification protocol using AEFFIWNA- functionalized resins include conducting *(i)* a shorter column loading at high product titer; *(ii)* two wash steps, respectively with PBS at pH 7.4 (5 CVs) and 0.25 M NaCl at pH 7.4; and *(iii)* elution with 1 M NaCl at pH 8.0. This process delivered yield of encapsidated genomes and cell-transducing virions respectively of 51.2% and 50.1% (corresponding to an eluate titer of 3.3·10^7^ IFU/mL), along with a HCP LRV ∼2.2 and hcDNA LRV ∼1.3.

**Figure 6.**
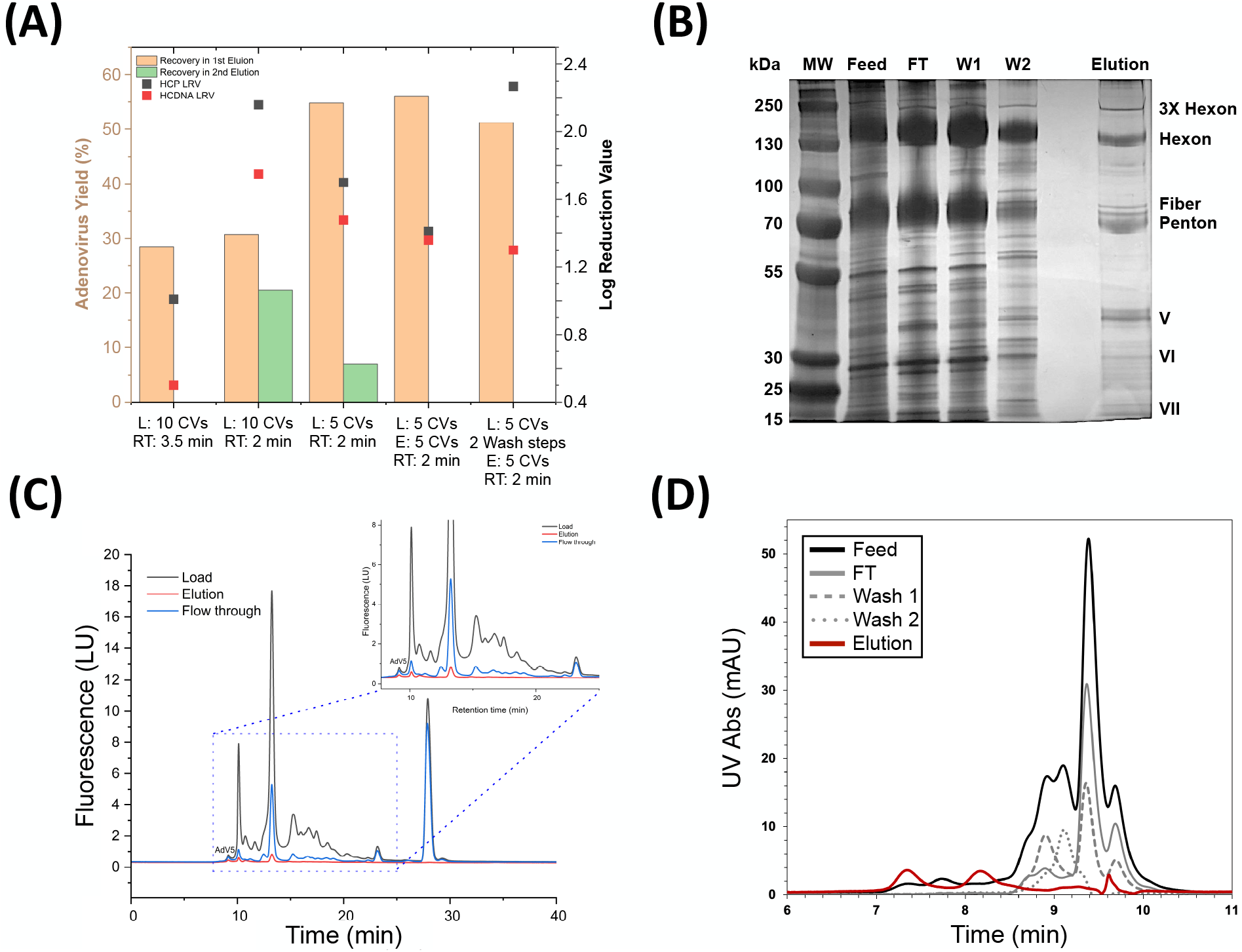
**(A)** Optimization of AdV5 purification process using AEFFIWNA-SulfoLink™ resins loaded with either 10 CVs of low AdV5 titer (∼5·10^9^ vg/mL) or 5 CVs of high AdV5 titer (∼2·10^10^ vg/mL) clarified HEK293 cell lysate at the RT of either 2 or 3.5 min, and washed in two steps prior to elution. The values of AdV5 yield were measured via RT-qPCR (encapsidated transgenes) and TCID50 assays (cell-transducing units); the reduction of HCPs and hcDNA were measured by analyzing the eluates and corresponding feedstocks using ELISA and PicoGreen™ dsDNA assay kits. **(B)** SDS-PAGE (non-reducing conditions) analysis of the chromatographic fractions produced by purifying AdV5 from a clarified HEK293 cell lysate using AEFFIWNA-SulfoLink™ resin; lane 1: molecular weight marker; lane 2: feedstock; lane 3: flow-through (5 CVs); lane 4: 1^st^ wash step at 0.15 M NaCl (PBS, 5 CVs); lane 5: 2^nd^ wash step at 0.25 M NaCl (5 CVs); lane 6: elution (5 CVs). SEC-HPLC analysis of the clarified HEK293 cell lysate and the flow-through, wash and elution fractions obtained by purifying AdV5 using AEFFIWNA-SulfoLink™ resin and analyzed using **(C)** a BioResolve SEC mAb HPLC column (200Å, 2.5 μm, 4.6 × 300 mm) and **(D)** a Bio SEC-5 HPLC column (2000 Å, 5 μm, 4.6 × 300 mm).

Selected chromatographic fractions were analyzed by SDS-PAGE and analytical size exclusion chromatography (SEC-HPLC), which provide an at-a-glance rendition of product concentration and contaminant removal. The electrophoretic analysis of the eluates in **Figure 6B** shows the presence of AdV capsid proteins, namely the monomeric and trimeric hexons (117 and 350 kDa; *note:* a study by Sweeney *et al*. attributes these bands to the AdV transgene) (Takahashi et al., 2006), pentons and fibers (75 and 70 kDa, respectively), and proteins V, VI, and VII (48 kDa, ∼24 kDa, and ∼18 kDa, respectively). The analytical chromatograms of the load and elution fractions obtained using two SEC columns with different exclusion limits provide further insight into the clearance of impurities in the flow-through and wash fractions (exclusion limit of 200 Å in **Figure 6C**) and the purity of the eluted AdVs (2000 Å in **Figure 6D**). The chromatograms of the load and flow through fractions in **Figure 6C** present a large amount of host cell contaminants (retention time ∼11 - 24 min); conversely, the eluate features two main peaks, corresponding to the whole AdV5 (∼9 min) and the hexon trimer (∼10 min). The AdV-associated peaks observed in the flow-through fraction suggest a slight overloading of the column, but they may also be caused by large, product-related contaminants (*e.g*., empty capsids and capsomers) whose size is above the exclusion limit of the SEC column. Therefore, a second SEC column, with larger exclusion limit, was adopted to better visualize the purity of eluted AdV5. The analytical chromatogram of the eluate in **Figure 6D** presents the peaks corresponding to the whole AdV5 (∼ 7 – 7.5 min) and the hexon trimer (∼8 min), and almost no residual contaminants (> 8.5 min); conversely, the latter form the majority of the species found in the chromatograms of the flow-through and wash fractions.

Collectively, these results demonstrate that the optimization of the virus titer in the lysate and of the chromatographic protocol grant high binding capacity and productivity of the affinity capture step along with excellent product yield (∼55% of encapsidated genomes and 50% of cell-transducing viral units) and quality in the eluate (residual titer of HCP ∼0.9 μg/mL and hcDNA ∼70 ng/mL).

### 3.4. Lifetime and performance of the affinity resin in a platform AdV purification process

The manufacturing of AdVs for oncolytic and vaccine applications currently relies on processes that employ anion-exchange resins or membranes operated in bind-and-elute mode for product capture and size-exclusion-mixed-mode chromatography in flow-through mode for product polishing. The adoption of anion-exchange chromatography requires the integration of multiple ultrafiltration/diafiltration (UF/DF) steps that adjust the conductivity and pH of the feed to each chromatography step, concentrate the product, and contribute to the removal of host cell contaminants (Bailey et al., 2014; Eglon et al., 2009; Turnbull et al., 2019). The proposed peptide-functionalized resins, with their excellent binding capacity and purification performance, are excellent candidates as affinity adsorbents for product capture in a new AdV purification platform. Furthermore, these adsorbents can process cell lysates without prior conditioning and release the product under the same elution conditions employed with anion-exchange adsorbents. We therefore assembled a platform AdV purification process comprising three steps: *(i)* clarification of the HEK293 cell lysate by depth filtration (*note:* the UF/DF step was omitted), *(ii)* product capture and purification by affinity chromatography in bind-and-elute mode using a peptide-functionalized affinity resin, and *(iii)* polishing by size-exclusion-mixed-mode chromatography using Capto™ Core 700 resin in flow-through mode (**Figure 7A**).

**Figure 7.**
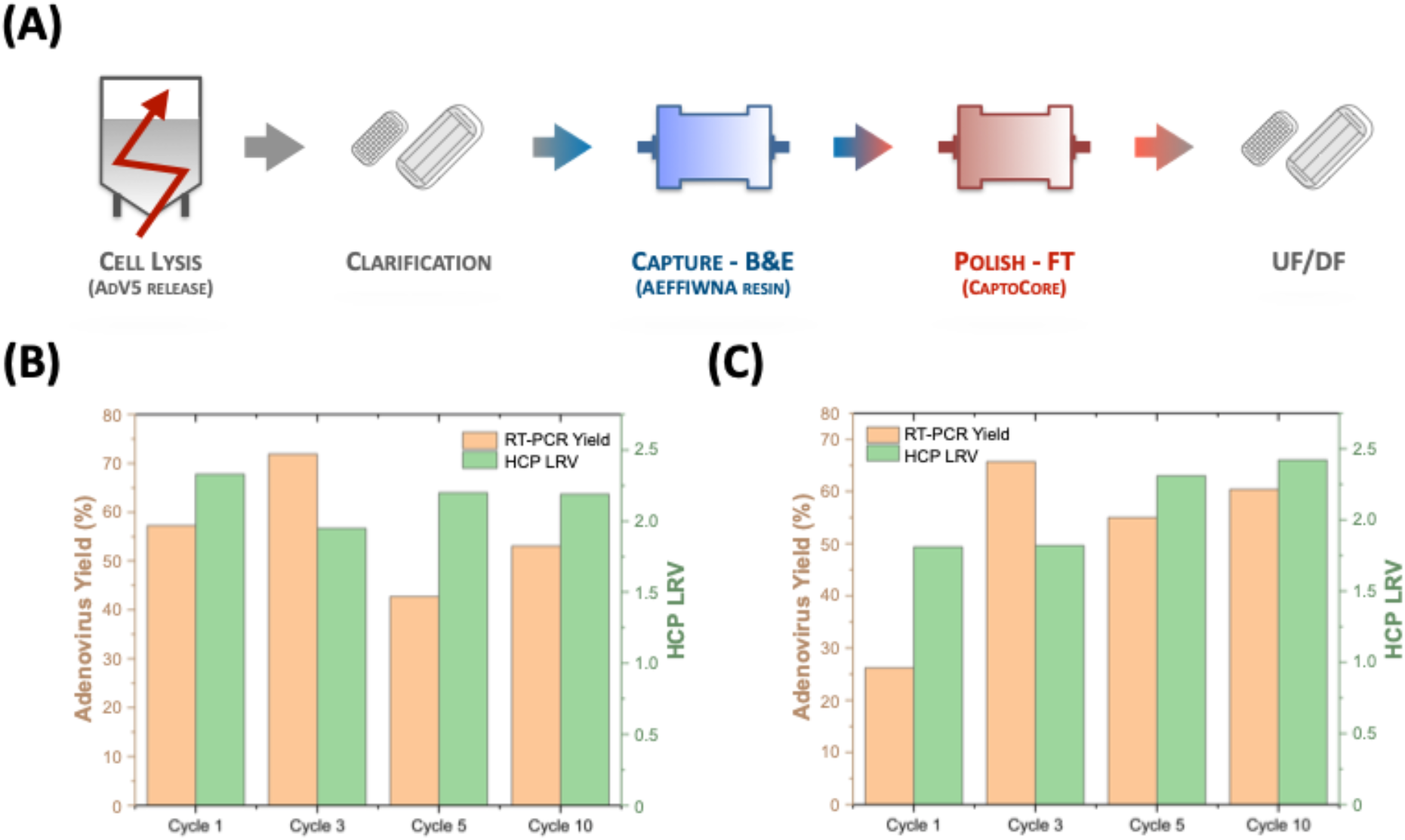
**(A)** AdV purification process comprising lysis of HEK293F cells, clarification via microfiltration, affinity-based capture using AEFFIWNA-functionalized resins in bind-and-elute mode, polishing with Capto™ Core 700 resin in flow-through mode. Performance of 10 cycles of AdV5 purification from a clarified HEK293 cell lysate (AdV5 titer ∼2·10^9^vg/mL; HCP titer ∼0.15 mg/mL) using **(B)** AEFFIWNA-Toyopearl® and **(C)** AEFFIWNA-SulfoLink™ resins with intermediate CIP using 10% v/v phosphoric acid. The values of AdV5 yield were measured via RT-qPCR (encapsidated transgenes).

The ability of an affinity resin to serve in an industrial process – especially one for manufacturing vaccines, whose price is expected to be lower than biological therapeutics – depends on its cost-effectiveness and lifetime. Linear peptide ligands, being produced synthetically, are significantly safer and more affordable than protein ligands produced recombinantly in bacterial systems, such as Protein A and the camelid ligands used for AAV purification. Specifically, the cost of goods of AEFFIWNA-functionalized resin is approximately $6.5-7.5K per liter when produced at beyond 10 liters scale. Regarding the resin lifetime, we sought to assay the stability of the resin to 10 cycles of AdV5 purification, each followed by a harsh cleaning-inplace (CIP): 10% v/v phosphoric acid (pH < 2) was used for AEFFIWNA-Toyopearl^®^, while 0.1 M glycine buffer at pH 2.0 was adopted for the CIP of AEFFIWNA-Sulfolink™ resin.

The resulting values of product yield and purity obtained with AEFFIWNA-functionalized resins across multiple purification cycles are collated in **Figure 7B** and **7C**, while the corresponding results obtained with TNDGPDYSSPLTGSG-Toyopearl^®^ are in **Figure S5**. All the corresponding chromatograms are collated in **Figure S6**. The affinity adsorbents, loaded to a ratio of ∼2.6·10^10^ vg per mL of resin (108% of DBC_10%_), did not show loss of binding capacity or selectivity across the successive purification cycles, affording average yields above 50%; the AEFFIWNA-functionalized resins provided HPC LRVs ∼ 2.0 - 2.3. The fluctuations in product yield and purity result from the inherent variability of cell lysates containing viral vectors and the analytical assays, but are consistent with the values recorded in prior work on the purification of AAVs and LVVs (Barbieri et al., 2024; Chu et al., 2023; Shastry et al., 2024a; Shastry et al., 2024b).

As the resin with the highest purification performance across multiple reuses, AEFFIWNA- SulfoLink™ resin was adopted as affinity resin for the capture step in the platform purification process outlined above. The values of step and global recovery and removal of host cell contaminants reported in **Table 2** demonstrate the effectiveness of the proposed technology for AdV purification. The global product yield of ∼55% aligns with the values obtained with processes that utilize anion-exchange resins and membranes (50 - 55%) (Anon, 2019; Peixoto et al., 2006). Most notably, the HCP titer was decreased ∼10,000-fold, from 0.1 mg/mL to 9.3 ng/mL; similarly, the hcDNA level was lowered ∼ 500 fold, from 5 μg/mL to 8.76 ng/mL, which is particularly remarkable since no DNase was utilized in the process.

**Table 2.**
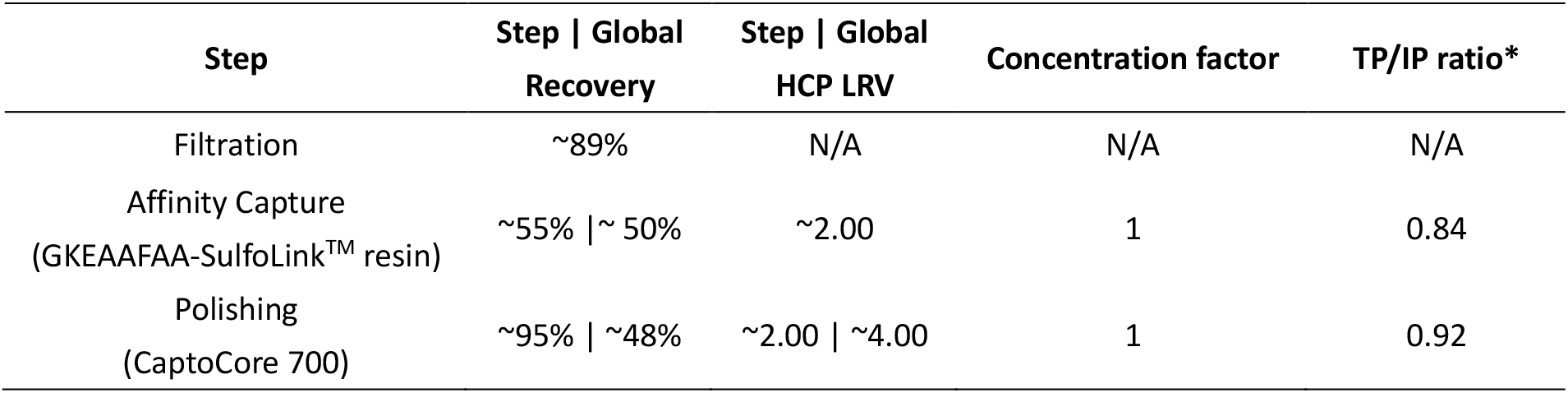
Values of AdV5 step and global recovery, removal of HEK293 HCPs and hcDNA, concentration factor, and total particles (TP) to infectious particles (IP) ratio obtained by purifying AdV5 from a HEK293 cell lysate (AdV5 titer ∼1·10^10^ vg/mL; HCP titer ∼0.15 mg/mL) using the process in **Figure 7A** (clarification: 0.45 um membrane sterile filtration; affinity capture: AEFFIWNA-SulfoLink™ resin).

| Step | Step \| Global<br>Recovery | Step \| Global<br>HCP LRV | Concentration factor | TP/IP ratio* |
| --- | --- | --- | --- | --- |
| Filtration | ~89% | N/A | N/A | N/A |
| Affinity Capture<br>(GKEAAFAA-SulfoLink™ resin) | ~55% \| ~ 50% | ~2.00 | 1 | 0.84 |
| Polishing<br>(CaptoCore 700) | ~95% \| ~48% | ~2.00 \| ~4.00 | 1 | 0.92 |

## 4. Conclusions

As the status of adenoviral vectors in oncolytic gene therapy and vaccination grows, modernizing the bioprocess technology for adenovirus purification is becoming of paramount importance. Particularly needed is the introduction of affinity adsorbents that increase the removal of process- and product-related contaminants far beyond the capabilities of ion-exchange adsorbents, while maintaining a high yield of encapsidated genomes and cell-transducing virions. In this context, hexon-targeting peptides demonstrate excellent potential as affinity ligands for adenovirus purification. The optimization of the adsorbent design and chromatographic operation grants high binding capacity at moderately high flow rates, the effective clearance of host cell proteins and nucleic acids, the removal of monomeric and multimeric capsid proteins, and the recovery of active virions under the same elution conditions utilized with anion-exchange media. Combined, these features translate in high productivity and product quality, and set the stage for peptide-functionalized resins to seamlessly replace traditional adsorbents in current adenovirus purification processes. Future endeavors will focus on the development of peptide sequences that resist alkaline cleaning-in-place and their evaluation as ligands for the serotype-agnostic purification of AdVs.

## Supporting information

AdV paper ESI

## Acknowledgements

The authors wish to acknowledge the funding provided by the Food and Drug Administration (Grant R01FD007481), the Novo Nordisk Foundation (AIM-Bio Grant NNF19SA0035474), the Triangle Universities Center for Advanced Studies Inc. (TUCASI), and the North Carolina Viral Vector Initiative in Research and Learning (NC-VVIRAL) at NC State University.

## Conflict of interest

The authors declare no conflict of interest.

## Notes

### Competing Interest Statement

The authors have declared no competing interest.

## References

1. Afkhami S, Yao Y, Xing Z. 2016. Methods and clinical development of adenovirus-vec-tored vaccines against mucosal pathogens. Mol Ther Methods Clin Dev 3:16030.

2. A Scalable Adenovirus Production Process from Cell Culture to Purified Bulk Product. 2019. https://www.genengnews.com/topics/bioprocessing/a-scalable-adenovirus-production-process-from-cell-culture-to-purified-bulk-product/.

3. FDA D.I.S.C.O. Burst Edition: FDA approval of Adstiladrin (nadofaragene firadenovec-vncg) for patients with high-risk Bacillus Calmette-Guérin unresponsive non-muscle invasive bladder cancer with carcinoma in situ with or without papillary tumors. 2022. https://www.fda.gov/drugs/resources-information-approved-drugs/fda-disco-burst-edition-fda-approval-adstiladrin-nadofaragene-firadenovec-vncg-patients-high-risk.

4. Bailey LJ, Sheehy KM, Hoey RJ, Schaefer ZP, Ura M, Kossiakoff AA. 2014. Applications for an engineered Protein-G variant with a pH controllable affinity to antibody fragments. J Immunol Methods 415:24–30.

5. Barbieri E, Mollica GN, Moore BD, Sripada SA, Shastry S, Kilgore RE, Loudermilk CM, Whitacre ZH, Kilgour KM, Wuestenhagen E, Aldinger A, Graalfs H, Rammo O, Schulte MM, Johnson TF, Daniele MA, Menegatti S. 2024. Peptide ligands targeting the vesicular stomatitis virus G (VSV‐G) protein for the affinity purification of lentivirus particles. Biotechnol Bioeng 121:618–639.

6. Barkats M, Bilang-Bleuel A, Buc-Caron MH, Castel-Barthe MN, Corti O, Finiels F, Horellou P, Revah F, Sabate O, Mallet J. 1998. Adenovirus in the brain: recent advances of gene therapy for neurodegenerative diseases. Prog Neurobiol 55:333–341.

7. Barton KN, Paielli D, Zhang Y, Koul S, Brown SL, Lu M, Seely J, Ho Kim J, Freytag SO. 2006. Second-Generation Replication-Competent Oncolytic Adenovirus Armed with Improved Suicide Genes and ADP Gene Demonstrates Greater Efficacy without Increased Toxicity. Molecular Therapy 13:347–356.

8. Brument N, Morenweiser R, Blouin V, Toublanc E, Raimbaud I, Chérel Y, Folliot S, Gaden F, Boulanger P, Kroner-Lux G, Moullier P, Rolling F, Salvetti A. 2002. A Versatile and Scalable Two-Step Ion-Exchange Chromatography Process for the Purification of Recombinant Adeno-associated Virus Serotypes-2 and -5. Molecular Therapy 6:678–686.

9. Burova E, Ioffe E. 2005. Chromatographic purification of recombinant adenoviral and adeno-associated viral vectors: methods and implications. Gene Ther 12:S5–S17.

10. Chu W, Prodromou R, Moore B, Elhanafi D, Kilgore R, Shastry S, Menegatti S. 2022. Development of peptide ligands for the purification of α-1 antitrypsin from cell culture fluids. J Chromatogr A 1679:463363.

11. Chu W, Shastry S, Barbieri E, Prodromou R, Greback‐Clarke P, Smith W, Moore B, Kilgore R, Cummings C, Pancorbo J, Gilleskie G, Daniele MA, Menegatti S. 2023. Peptide ligands for the affinity purification of adeno‐associated viruses from HEK 293 cell lysates. Biotechnol Bioeng 120:2283–2300.

12. Colbert L, Jia Y, Sharma A, Hu J, Xu Z, Suzman DL, Das A, Bross P, Kluetz PG, Fashoyin-Aje LA. 2025. FDA Approval Summary: Nadofaragene Firadenovec-vncg for Bacillus Calmette–Guérin–Unresponsive Non–Muscle-Invasive Bladder Cancer. Clinical Cancer Research:OF1–OF4.

13. Dietl S, Kiefer F, Binder S, Walther P, Sobek H, Mizaikoff B. 2023. An efficient capture strategy for the purification of human adenovirus type 5 from cell lysates. J Biotechnol 361:49– 56.

14. Eglon MN, Duffy AM, O’Brien T, Strappe PM. 2009. Purification of adenoviral vectors by combined anion exchange and gel filtration chromatography. J Gene Med 11:978–989.

15. Ehrke-Schulz E, Zhang W, Schiwon M, Bergmann T, Solanki M, Liu J, Boehme P, Leitner T, Ehrhardt A. 2016. Cloning and Large-Scale Production of High-Capacity Adenoviral Vectors Based on the Human Adenovirus Type 5. Journal of Visualized Experiments.

16. Gao J, Zhang W, Ehrhardt A. 2020. Expanding the Spectrum of Adenoviral Vectors for Cancer Therapy. Cancers (Basel) 12:1139.

17. Gast M, Sobek H, Mizaikoff B. 2019. Nanoparticle Tracking of Adenovirus by Light Scattering and Fluorescence Detection. Hum Gene Ther Methods 30:235–244.

18. Gast M, Wondany F, Raabe B, Michaelis J, Sobek H, Mizaikoff B. 2020. Use of Super-Resolution Optical Microscopy To Reveal Direct Virus Binding at Hybrid Core–Shell Matrixes. Anal Chem 92:3050–3057.

19. Inger Salomonsson. 2022. 2-Step Purification of Adenovirus using Ion Exchange chromatography. https://www.cytivalifesciences.com/en/us/news-center/adenovirus-purification-10001?srsltid=AfmBOoq4g1TaaGfcEzKzjDGtiHkeYiqPQ3vzXSg1_9PIFFt8c8gtVBdq.

20. Iyer P, Ostrove JM, Vacante D. 1999. Comparison of manufacturing techniques for adenovirus production. Cytotechnology 30:169–172.

21. Kaludov N, Handelman B, Chiorini JA. 2002. Scalable Purification of Adeno-Associated Virus Type 2, 4, or 5 Using Ion-Exchange Chromatography. Hum Gene Ther 13:1235–1243.

22. Lee CS, Bishop ES, Zhang R, Yu X, Farina EM, Yan S, Zhao C, Zeng Z, Shu Y, Wu X, Lei J, Li Y, Zhang W, Yang C, Wu K, Wu Y, Ho S, Athiviraham A, Lee MJ, Wolf JM, Reid RR, He T-C. 2017. Adenovirus-mediated gene delivery: Potential applications for gene and cell-based therapies in the new era of personalized medicine. Genes Dis 4:43–63.

23. Nadeau I, Kamen A. 2003. Production of adenovirus vector for gene therapy. Biotechnol Adv 20:475–489.

24. Nokisalmi P, Pesonen S, Escutenaire S, Särkioja M, Raki M, Cerullo V, Laasonen L, Alemany R, Rojas J, Cascallo M, Guse K, Rajecki M, Kangasniemi L, Haavisto E, Karioja-Kallio A, Hannuksela P, Oksanen M, Kanerva A, Joensuu T, Ahtiainen L, Hemminki A. 2010. Oncolytic Adenovirus ICOVIR-7 in Patients with Advanced and Refractory Solid Tumors. Clinical Cancer Research 16:3035–3043.

25. Peixoto C, Ferreira TB, Carrondo MJT, Cruz PE, Alves PM. 2006. Purification of adenoviral vectors using expanded bed chromatography. J Virol Methods 132:121–126.

26. Ricobaraza A, Gonzalez-Aparicio M, Mora-Jimenez L, Lumbreras S, Hernandez-Alcoceba R. 2020. High-Capacity Adenoviral Vectors: Expanding the Scope of Gene Therapy. Int J Mol Sci 21:3643.

27. Sakurai F, Tachibana M, Mizuguchi H. 2022. Adenovirus vector-based vaccine for infectious diseases. Drug Metab Pharmacokinet 42:100432.

28. Salauddin Md, Saha S, Hossain MdG, Okuda K, Shimada M. 2024. Clinical Application of Adenovirus (AdV): A Comprehensive Review. Viruses 16:1094.

29. Shabram PW, Giroux DD, Goudreau AM, Gregory RJ, Horn MT, Huyghe BG, Liu X, Nunnally MH, Sugarman BJ, Sutjipto S. 1997. Analytical Anion-Exchange HPLC of Recombinant Type-5 Adenoviral Particles. Hum Gene Ther 8:453–465.

30. Shastry S, Barbieri E, Minzoni A, Chu W, Johnson S, Stoops M, Pancorbo J, Gilleskie G, Ritola K, Crapanzano MS, Daniele MA, Menegatti S. 2024a. Serotype-agnostic affinity purification of adeno-associated virus (AAV) via peptide-functionalized chromatographic resins. J Chromatogr A 1734:465320.

31. Shastry S, Chu W, Barbieri E, Greback‐Clarke P, Smith WK, Cummings C, Minzoni A, Pancorbo J, Gilleskie G, Ritola K, Daniele MA, Johnson TF, Menegatti S. 2024b. Rational design and experimental evaluation of peptide ligands for the purification of adeno‐associated viruses via affinity chromatography. Biotechnol J 19.

32. Sorensen MR, Holst PJ, Pircher H, Christensen JP, Thomsen AR. 2009. Vaccination with an adenoviral vector encoding the tumor antigen directly linked to invariant chain induces potent CD4 ^+^ T‐cell‐independent CD8 ^+^ T‐cell‐mediated tumor control. Eur J Immunol 39:2725– 2736.

33. Syyam A, Nawaz A, Ijaz A, Sajjad U, Fazil A, Irfan S, Muzaffar A, Shahid M, Idrees M, Malik K, Afzal S. 2022. Adenovirus Vector System: Construction, History and Therapeutic Applications. Biotechniques 73:297–305.

34. Takahashi E, Cohen SL, Tsai PK, Sweeney JA. 2006. Quantitation of adenovirus type 5 empty capsids. Anal Biochem 349:208–217.

35. Tasca F, Wang Q, Gonçalves MAFV. 2020. Adenoviral Vectors Meet Gene Editing: A Rising Partnership for the Genomic Engineering of Human Stem Cells and Their Progeny. Cells 9:953.

36. Thacker EE, Timares L, Matthews QL. 2009. Strategies to overcome host immunity to adenovirus vectors in vaccine development. Expert Rev Vaccines 8:761–777.

37. Turnbull JP. Development of a High Recovery Adenovirus Purification Process Using Anion Exchange Nanofibers.

38. Turnbull J, Wright B, Green NK, Tarrant R, Roberts I, Hardick O, Bracewell DG. 2019. Adenovirus 5 recovery using nanofiber ion‐exchange adsorbents. Biotechnol Bioeng 116:1698–1709.

39. Ugai H, Yamasaki T, Hirose M, Inabe K, Kujime Y, Terashima M, Liu B, Tang H, Zhao M, Murata T, Kimura M, Pan J, Obata Y, Hamada H, Yokoyama KK. 2005. Purification of infectious adenovirus in two hours by ultracentrifugation and tangential flow filtration. Biochem Biophys Res Commun 331:1053–1060.

40. Wang Z-X, Bian H-B, Yang J-S, De W, Ji X-H. 2009. Adenovirus-mediated suicide gene therapy under the control of Cox-2 promoter for colorectal cancer. Cancer Biol Ther 8:1480– 1488.

41. Watanabe M, Nishikawaji Y, Kawakami H, Kosai K. 2021. Adenovirus Biology, Recombinant Adenovirus, and Adenovirus Usage in Gene Therapy. Viruses 13:2502.

42. Wiethoff CM, Wodrich H, Gerace L, Nemerow GR. 2005. Adenovirus Protein VI Mediates Membrane Disruption following Capsid Disassembly. J Virol 79:1992–2000.

43. Wold W, Toth K. 2014. Adenovirus Vectors for Gene Therapy, Vaccination and Cancer Gene Therapy. Curr Gene Ther 13:421–433.

44. Wu S, Huang GY, Liu J. 2002. Production of Retrovirus and Adenovirus Vectors for Gene Therapy: A Comparative Study Using Microcarrier and Stationary Cell Culture. Biotechnol Prog 18:617–622.

45. Wu Y, Barbieri E, Kilgore RE, Moore BD, Chu W, Mollica GN, Daniele MA, Menegatti S. 2024. Peptide ligands for the affinity purification of adenovirus from HEK293 and vero cell lysates. J Chromatogr A 1736:465396.

46. Yang H. 2013. Establishing Acceptable Limits of Residual DNA. PDA J Pharm Sci Technol 67:155–163.

