## Supplementary material for "Integrating affinity chromatography in the platform process for Adenovirus purification": AdV paper ESI

**S.1. Measuring the cell-transduction activity of purified AdV5.** To evaluate the activity of the AdV5 eluted from peptide-functionalized resins, we performed transduction and transfection assays. The AdVs utilized in this study encapsidate a transgene encoding for green fluorescent protein (GFP), which enables in principle the quantification of the cell-transducing viral particles by fluorescence flow cytometry. However, during the incubation time required to complete a comprehensive transduction assay, the host cells develop cytopathic effects leading to significant cell death and a decline in cell growth. Consequently, despite observing a strong green fluorescent signal during flow cytometry, the limited cell counts prevented the derivation of quantitative metrics. Nonetheless, the qualitative observation of the pronounced fluorescent signal documented the transduction capability of the purified AdV5. In lieu of the transduction assay, a label-free cytopathic effect (CPE) assay was employed to assess the tissue culture infectious dose (TCID50) of AdV5 in the clarified lysates and the affinity eluates. A microscopic illustration of the cytopathic effect is presented in **Figure S1**. Because AdVs cannot be propagated in Vero cells without introducing exogenous E1, HEK293 adherent cells were utilized for all TCID50 assays (see methods in *Section 2.8*). Aligning with the values of yield of encapsidated transgenes measured by RT-qPCR, the transfection assays returned values of yield from HEK293 and Vero cell lysates of 46.4% and 43.1% using AEFFIWNA-Toyopearl^®^ resin, and 50.1% and 43.6% using TNDGPDYSSPLTGSG-Toyopearl^®^ resin (**Table S1**). These results confirm that the selected peptide ligands yield a product with high purity and activity under mild conditions, validating the ligand development strategy presented in prior work.

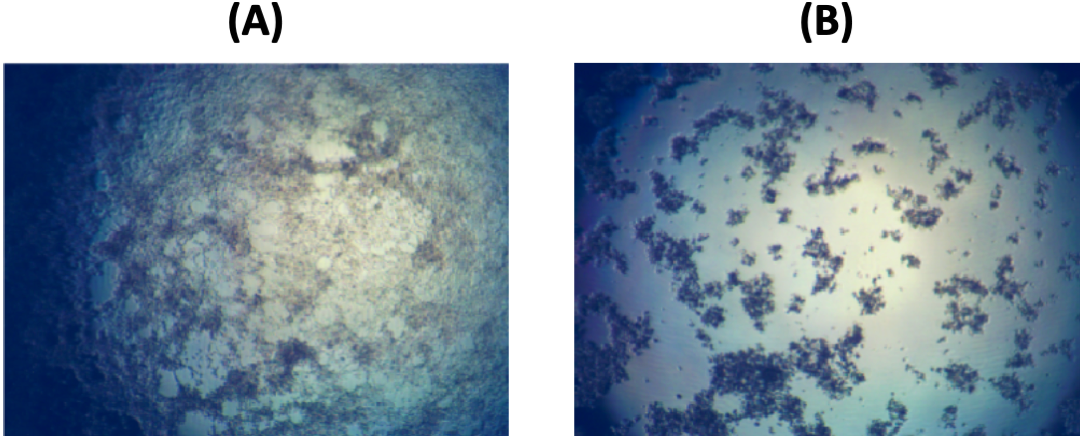

***Figure S1. (A)*** *TCID50 control well;* ***(B)*** *cells displaying cytopathic effects after transfection by AdV5 purified from a clarified HEK293 cell lysate using AEFFIWNA-Toyopearl^®^ resin.*

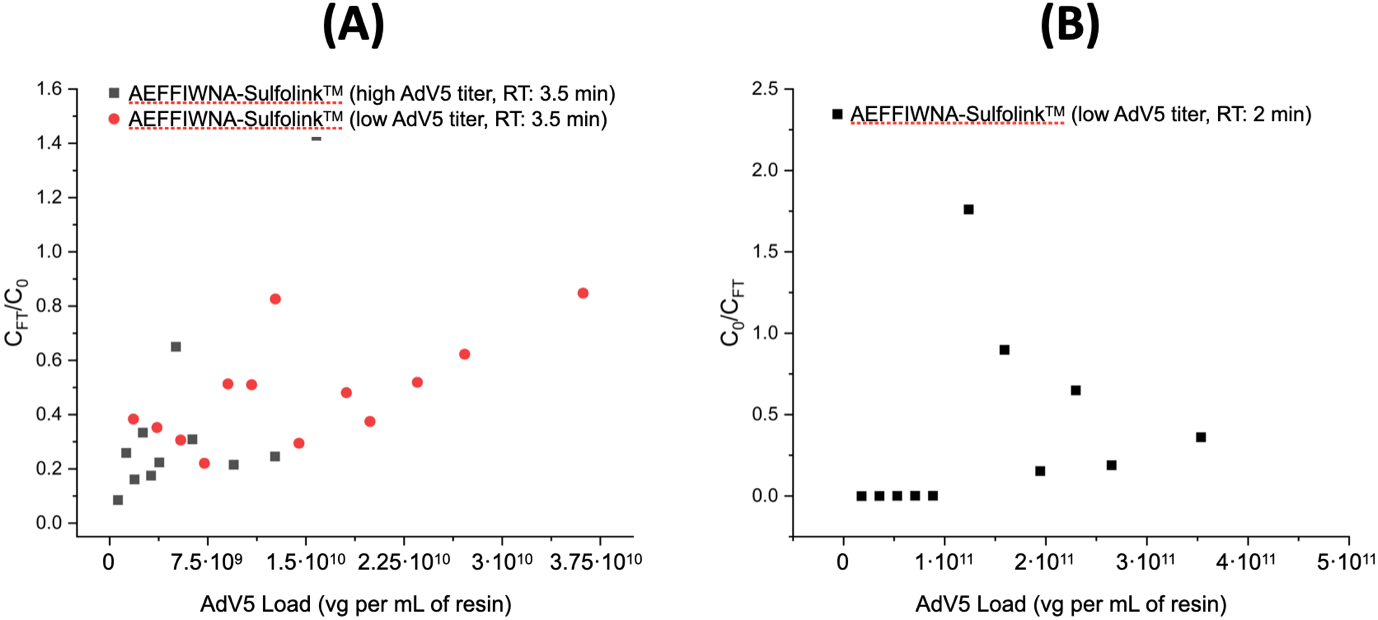

***Figure S2.*** *Breakthrough curves of AdV5 encapsidated transgenes obtained by loading clarified HEK293 cell lysate (AdV5 titer ~6.3*10^8^ or ~1.0*·*10^10^ vg/mL; HCP titer ~0.15 mg/mL) on AEFFIWNA-SulfoLink^TM^ resin loaded at the RT of 3.5 min.*

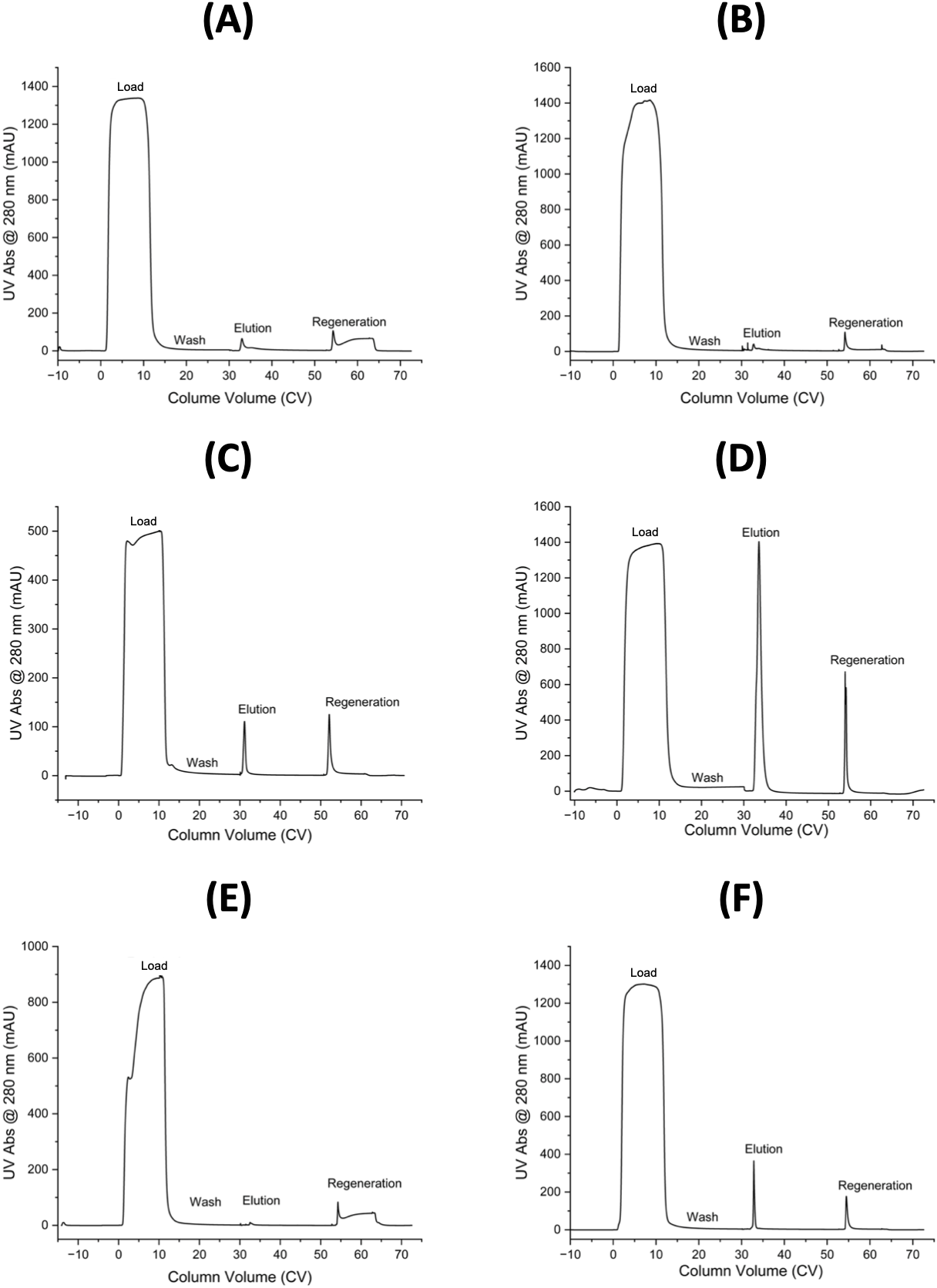

***Figure S3.*** *Chromatograms* *of AdV5 purification from a clarified HEK293 cell lysate (AdV5 titer ~10^9^ vg/mL; HCP titer ~0.15 mg/mL) using* ***(A)*** *AEFFIWNA-Toyopearl® resin,* ***(B)*** *AEFFIWNA-Poros^TM^ resin,* ***(C)*** *AEFFIWNA-SulfoLink^TM^ resin,* ***(D)*** *TNDGPDYSSPLTGSG-Toyopearl® resin,* ***(E)*** *TNDGPDYSSPLTGSG-Poros^TM^ resin, and* ***(F)*** *TNDGPDYSSPLTGSG-SulfoLink^TM^ resin.*

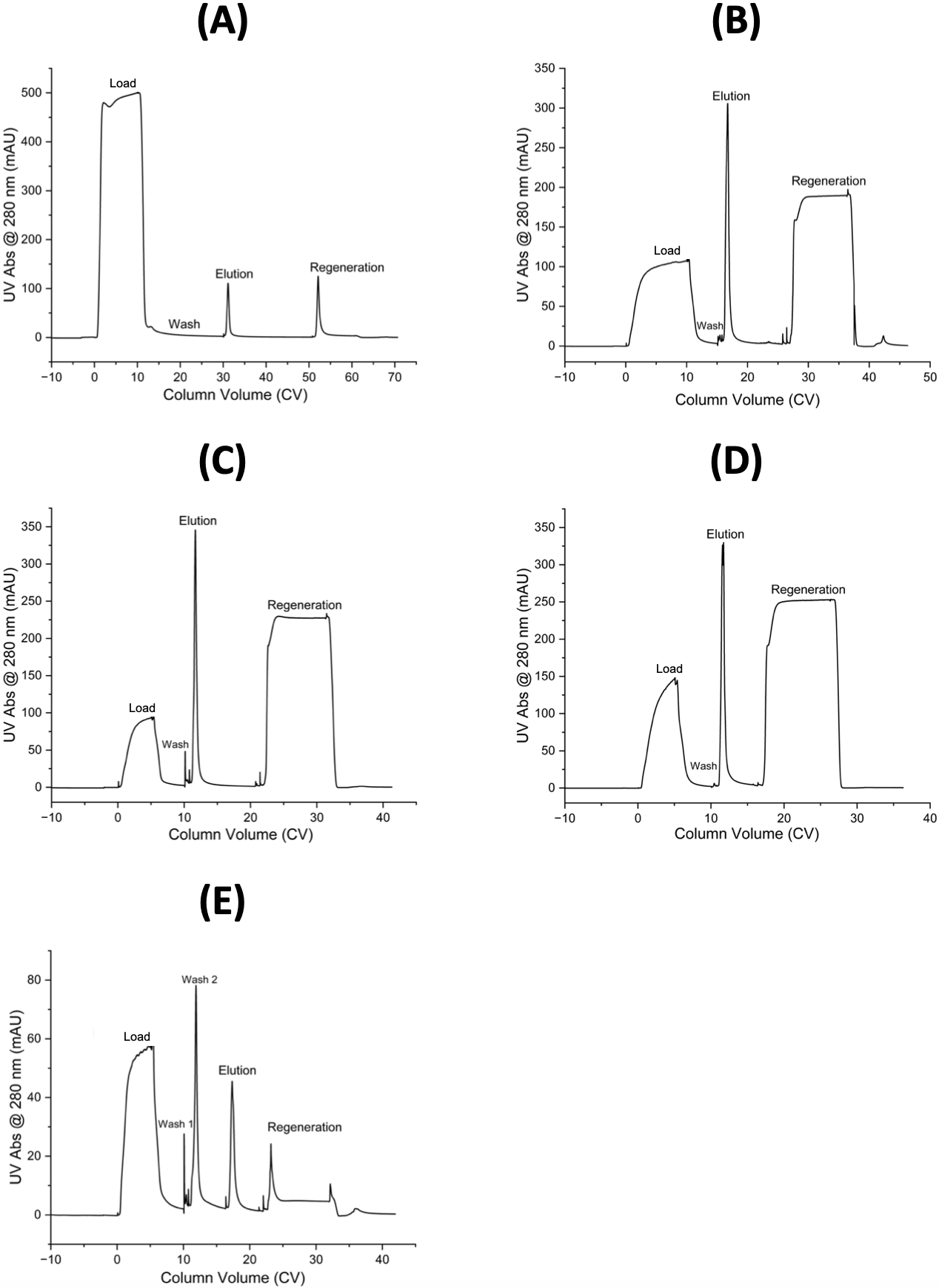

***Figure S4.*** *Chromatograms* *of AdV5 purification using AEFFIWNA-SulfoLink^TM^ resin loaded with 10 CVs of low AdV5 titer (~5·10^9^ vg/mL) at the RT of* ***(A)*** *3.5 min or* ***(B)*** *2 min. Chromatograms* *of AdV5 purification using AEFFIWNA-SulfoLink^TM^ resin loaded with 5 CVs of high AdV5 titer (~2·10^10^ vg/mL) clarified HEK293 cell lysate and eluted with* ***(C)*** *10 CVs or* ***(D)*** *5 CVs.* ***(E)*** *Chromatogram* *of AdV5 purification using AEFFIWNA-SulfoLink^TM^ resin loaded with 5 CVs of high AdV5 titer (~2·10^10^ vg/mL) clarified HEK293 cell lysate, exposed to two wash steps, and eluted with 5 CVs.*

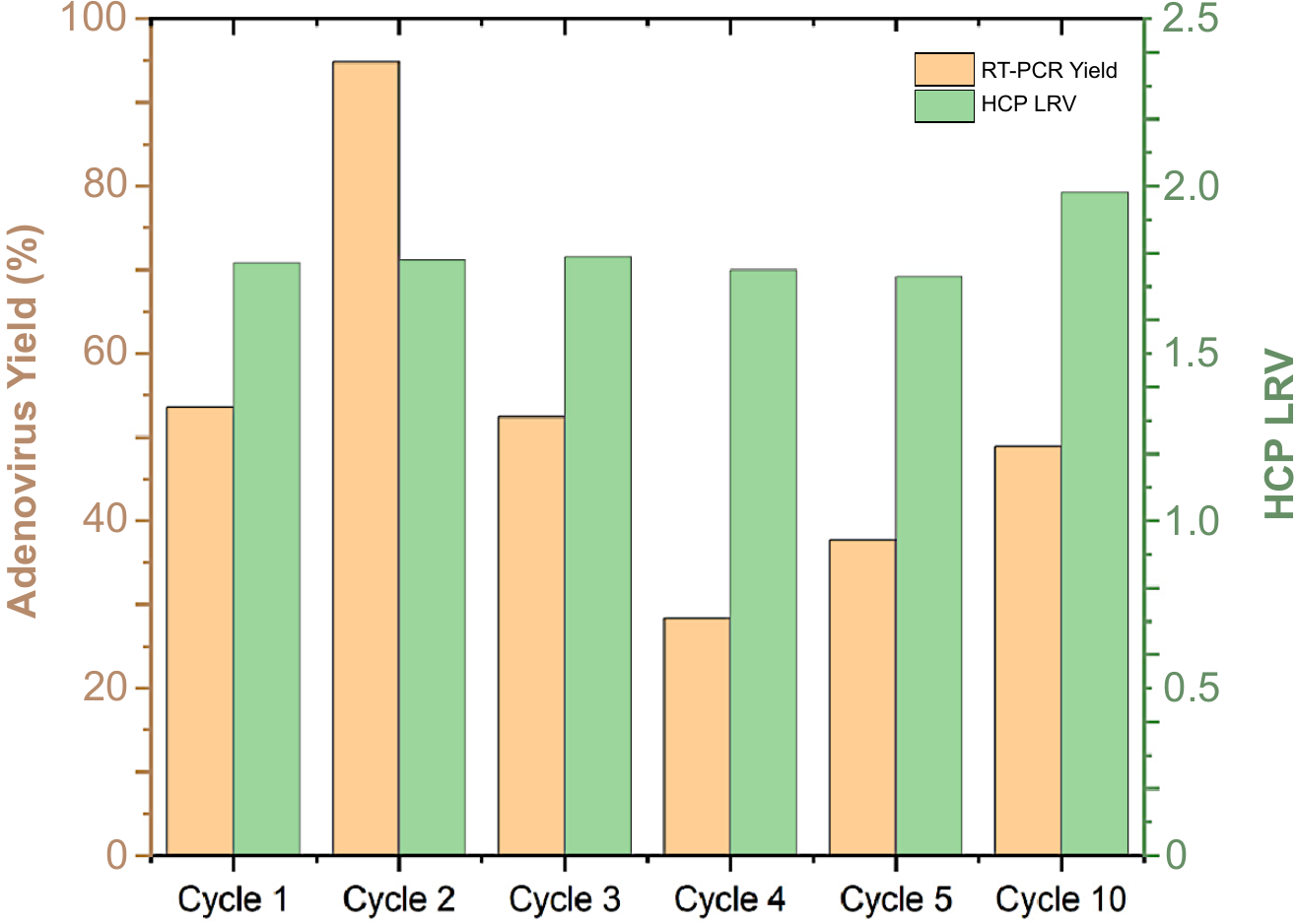

***Figure S5.*** *Performance of 10 cycles of AdV5 purification from a clarified HEK293 cell lysate (AdV5 titer ~2.0·10^8^ vg/mL; HCP titer ~0.15 mg/mL) using* *TNDGPDYSSPLTGSG-Toyopearl^®^ resin with intermediate CIP using 10% v/v phosphoric acid. The values of AdV5 yield were measured via RT-qPCR (encapsidated transgenes); the reduction of HCPs was measured by analyzing the eluates and corresponding feedstocks using ELISA kits.*

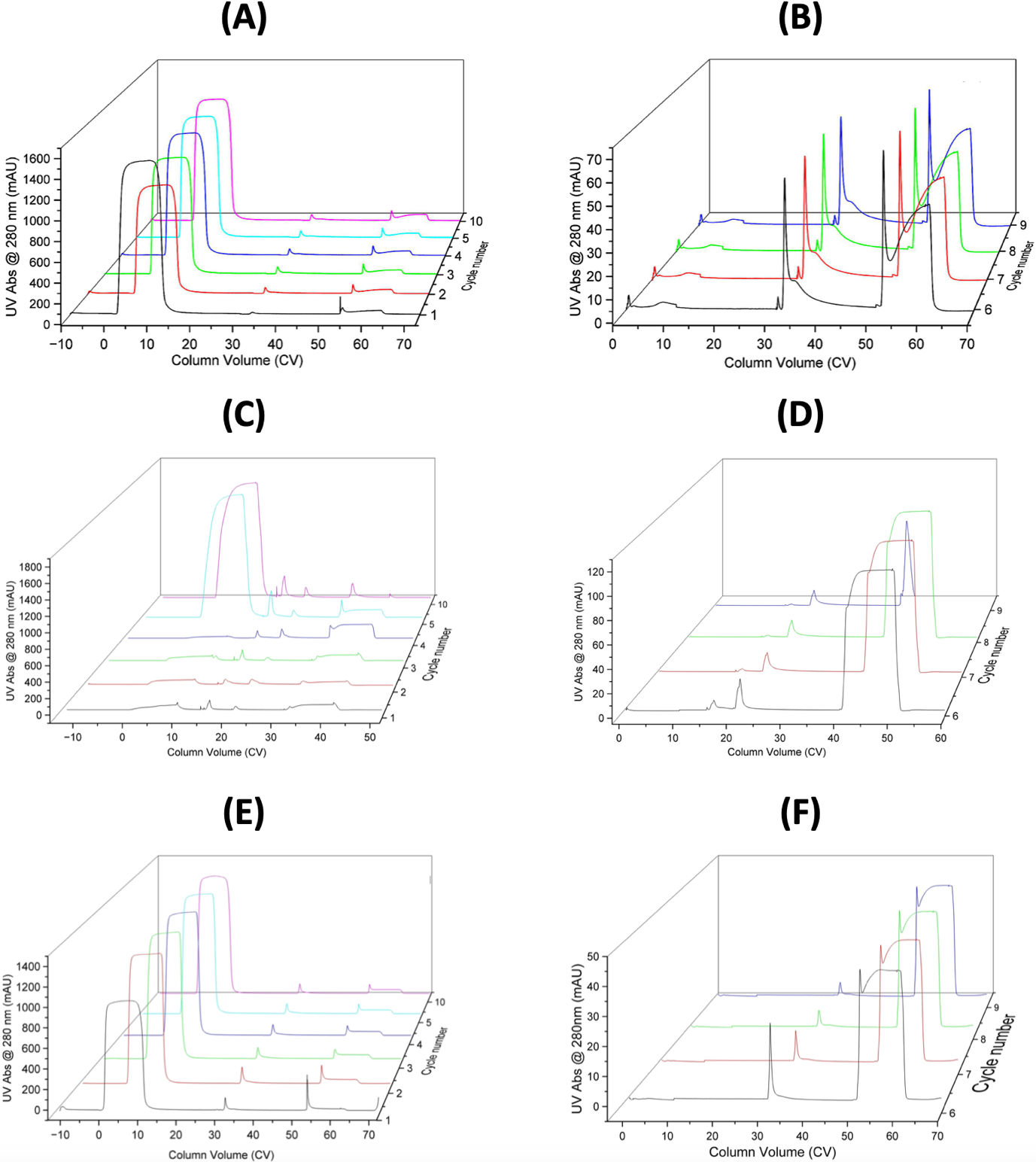

***Figure S6.*** *Chromatogram of* ***(A)*** *five cycles of AdV5 purification from a clarified HEK293 cell lysate (AdV5 titer ~2·10^9^vg/mL; HCP titer ~0.15 mg/mL) using AEFFIWNA-Toyopearl® resin and* ***(B)*** *five cycles of blank loading on AEFFIWNA-Toyopearl® resin. Chromatogram of* ***(C)*** *five cycles of AdV5 purification from a clarified HEK293 cell lysate (AdV5 titer ~2·10^9^vg/mL; HCP titer ~0.15 mg/mL) using AEFFIWNA-SulfoLink^TM^ resin and* ***(D)*** *five cycles of blank loading on AEFFIWNA-SulfoLink^TM^ resin. Chromatogram of* ***(C)*** *five cycles of AdV5 purification from a clarified HEK293 cell lysate (AdV5 titer ~2·10^9^vg/mL; HCP titer ~0.15 mg/mL) using TNDGPDYSSPLTGSG-Toyopearl^®^ resin and* ***(D)*** *five cycles of blank loading on TNDGPDYSSPLTGSG-Toyopearl^®^ resin.*

***Table*** ***S1.*** *Purification of AdV5 from clarified HEK293 and Vero cell lysates using AEFFIWNA-Toyopearl*® *and TNDGPYSSPLTGSG-Toyopearl*® *resins. The lysates were loaded at the residence time of 3.5 min to a ratio of 5·10*^9^ *vg per mL of resin. The bound AdV5 were eluted in 20 mM Tris buffer with 1 M NaCl at pH 8.0. The values of AdV5 titer were measured via RT-qPCR (encapsidated transgenes) and TCID50 assays (cell-transducing units); the HCP and hcDNA titers were measured by analyzing the eluates and corresponding feedstocks using ELISA and PicoGreen™ dsDNA assay kits.*

| **Peptide** | **Feedstock** | | | | **Eluate** | | | | | | |
| --- | --- | --- | --- | --- | --- | --- | --- | --- | --- | --- | --- |
|  | **Cell Source** | **AdV titer**  **(vg/mL)** | **hcDNA titer**  **(μg/mL)** | **HCP titer**  **(mg/mL)** | **AdV titer**  **(vg/mL)** | **hcDNA titer**  **(μg/mL)** | **HCP titer**  **(μg/mL)** | **Gene Yield**  **(RT-PCR)** | **TU Yield** | **HCP LRV** | **hcDNA LRV** |
| AEFFIWNA | HEK293 | 1.5·10^9^ | 12.3 | 0.35 | 3.1·10^8^ | 0.118 | 1.4 | 40.8% | 46.4% | 2.07 | 1.81 |
|  | Vero | 4.0·10^8^ | 5.3 | 0.14 | 1.1·10^8^ | 0.149 | 0.85 | 53.2% | 43.1% | 1.92 | 1.34 |
| TNDGPDYSSPLTGSG | HEK293 | 4.4·10^8^ | 21.8 | 0.16 | 1.0·10^8^ | 0.43 | 1.54 | 50.3% | 50.1% | 1.72 | 1.41 |
|  | Vero | 2.0·10^9^ | 6.8 | 0.08 | 5.6·10^8^ | 0.15 | 1.3 | 55.5% | 43.6% | 1.60 | 1.36 |

***Table S2.*** *Purification of AdV5 from clarified a HEK293 cell lysate using different chromatographic resins functionalized with peptide ligands AEFFIWNA and TNDGPYSSPLTGSG. The lysates were loaded at the residence time of 3.5 min to a ratio of 5·10^10^ vg per mL of resin. The bound AdV5 were eluted in 20 mM Tris buffer with 1 M NaCl at pH 8.0. The values of AdV5 titer were measured via RT-qPCR (encapsidated transgenes) and TCID50 assays (cell-transducing units); the HCP and hcDNA titers were measured by analyzing the eluates and corresponding feedstocks using ELISA and PicoGreen™ dsDNA assay kits.*

| **Resin** | **Feedstock** | | | **Eluate** | | | | | | | |
| --- | --- | --- | --- | --- | --- | --- | --- | --- | --- | --- | --- |
|  | **AdV titer**  **(vg/mL)** | **HCP titer**  **(μg/mL)** | **hcDNA titer**  **(μg/mL)** | | **AdV titer**  **(vg/mL)** | **HCP titer**  **(μg/mL)** | **hcDNA titer**  **(μg/mL)** | **Gene Yield**  **(RT-PCR)** | **TU Yield** | **HCP LRV** | **hcDNA LRV** |
| AEFFIWNA-Toyopearl | 1.5·10^9^ | 0.353 | 12.3 | | 3.1·10^8^ | 1.35 | 0.118 | 40.8% | 46.4% | 2.07 | 1.81 |
| AEFFIWNA-Poros | 9·10^8^ | 0.216 | 2.9 | | 2.0·10^8^ | 2.05 | 0.203 | 43.2% | 32.1% | 1.72 | 0.85 |
| AEFFIWNAC-Sulfolink | 5.6·10^10^ | 0.551 | 2.8 | | 8.0·10^9^ | 26.9 | 0.457 | 28.5% | 33.3% | 1.01 | 0.50 |
| TNDGPDYSSPLTGSG-Toyopearl | 4.4·10^8^ | 0.160 | 21.8 | | 1.1·10^8^ | 1.54 | 0.43 | 50.3% | 50.1% | 1.72 | 1.41 |
| TNDGPDYSSPLTGSG-Poros | 2.2·10^8^ | 0.067 | 2.3 | | 2.4·10^7^ | 1.06 | 0.036 | 21.8% | 12.6% | 1.50 | 1.50 |
| TNDGPDYSSPLTGSGC-Sulfolink | 1.2·10^10^ | 0.044 | 2.3 | | 4.2·10^8^ | 0.153 | 0.103 | 7.0% | < 1% | 2.15 | 1.05 |
